# Variability in scRNA-Seq analysis is mitigated by regularized or supervised approaches

**DOI:** 10.1101/2021.02.15.431268

**Authors:** Arda Durmaz, Jacob G. Scott

## Abstract

Transcriptional dynamics of evolutionary processes through time are highly complex and require single-cell resolution datasets. This is especially important in cancer during the evolution of resistance, where stochasticity can lead to selection for divergent transcriptional mechanisms. Statistical methods developed to address various questions in single-cell datasets are prone to variability and require careful adjustments of multiple parameter space. To assess the impact of this variation, we utilized commonly used single-cell RNA-Seq analysis tools in a combinatorial fashion to evaluate how repeatable the results are when different methods are combined. In the context of clustering and trajectory estimation, we benchmark the combinatorial space and highlight ares and methods that are sensitive to parameter changes. We have observed that utilizing temporal information in a supervised framework or regularization in latent modeling reduces variability leading to improved overlap when different parameters/methods are used. We hope that future studies can benefit from the results presented here as use of scRNA-Seq analysis tools as out of the box is becoming a standard approach in cancer research.

## Introduction

Intra-tumor heterogeneity has recently become a central focus of cancer research secondary to the limited response of patients to targeted therapies. These failures are driven by Darwinian evolution, by heritable variation and selection through time. One source for the subsequent intra-tumor heterogeneity is the variation driven by stochasticity in transcriptional activity modulated by epigenetic processes^[1,2]^. This change in overall composition is further modulated by the selective advantage of pre-existing resistant cells or clonal expansion of drug tolerant cells mediated by complex interactions between cells and the microenvironment^[3,4,5,6]^. Although previous efforts have made significant progress in understanding the complex cancer dynamics using bulk sequencing data, single-cell sequencing methods have allowed for novel insights by probing this heterogeneity directly – including during temporally varying processes. For instance, Lee *et al*. identified transcriptional heterogeneity as one of the key factors for promoting clonal expansion of drug-tolerant sub-population leading to the evolution of resistance^[7]^. Similarly, Kim *et al*. identified distinct sub-populations resistant to treatment in lung adenocarcinoma patients using single-cell RNA-Seq,^[8]^ Furthermore, relatively recently, single-cell sequencing coupled with mathematical models allowed for investigation of Darwinian dynamics, specifically treatment-induced selection pressure and transcriptional stochasticity at the single cell level^[9,10,11]^.

Investigating transcriptional regulation, single-cell sequencing also paved the way for pseudotime/trajectory estimation (PTE) to delineate temporal dynamics during differentiation and resistance evolution. Specifically, PTE aims to find low-dimensional proxy for the underlying transcriptional activity accounting for the temporal information. However, due to the stochasticity inherent in evolution, PTE poses additional challenges where replicate experiments can show divergent dynamics leading to evolution of distinct resistance mechanisms^[12,13]^. For instance, during multipotent progenitor trophoblast differentiation, stages of organization (endpoints) are clearly defined based on morphological characteristics hence we can reliably deduce functional mechanisms through time^[14]^. However, as we show in detail below, using the same analysis methods with slight differences in pre-processing parameters (number of genes expressed), can result in very different PTE orderings of cells in the setting of the evolution of resistance leading to increased diversity of identified mechanisms. Analysis of single-cell data is further complicated by the technical noise in library preparation strategies due to capture efficiencies at both cell (empty/multi-cell droplets) and transcript level.

In order to alleviate some of the issues with single-cell analysis, various analysis methods aim for robust imputation, outlier detection and quantification of gene expression. For instance, previous studies utilized imputation methods to reduce the effects of zero-counts due to dropouts in RNA-Seq datasets^[15,16]^. However, the increased number of available tools, and continued proliferation of them also requires careful selection of methods and associated parameters which can result in significant differences. This issue has been partially addressed before. Specifically, two comprehensive combinatorial evaluation studies have been conducted in order to evaluate different analysis workflows^[17,18]^. Tian *et al*. using cell-mixture experiments showed relatively good correlations between ground-truth and estimated trajectories using Slingshot or DDRTree^[18]^. Similarly, Saelens *et al*. showed improved performance for these methods using topological similarity metrics^[17]^. While illuminating, a major limitation of these studies is that the methods are applied on non-cancer (embryologic differentiation) processes or cancer cells in relatively homogeneous settings (without selection pressure). For instance, mixture experiments conducted by Tian *et al*. are limited to linear trajectories. In contrast, evolution under selection pressure can result in increased variability and non-linear patterns of transcriptional change^[19,20,21]^. As most tumors do not grow in these conditions, it is crucial to evaluate the available methods under selection pressure with temporal information as well. For this purpose, in this manuscript we report a benchmarking study in which we evaluate the available methods in a combinatorial fashion similar to Tian *et al*. and Saelens *et al*. focusing on the repeatability of PTEs. We hope that by evaluating the scRNA-Seq methods rigorously for settings applicable to the evolution of resistance in cancer, we will enable more robust and reproducible application of single cell sequencing technologies and experimental designs for future studies.

## Methods

We have focused on the repeatability of identified PTEs, and are hence utilizing methods widely adopted in the community. Additionally we utilize a neural network approach for dimension reduction to evaluate whether more complex models show any advantage when high-throughput single-cell datasets are used. Since neural networks have been extensively utilized for wide variety of problems in the form of autoencoders^[22]^ and relatively recently stochastic alternatives have been used for -omics datasets as well^[23,24,25,26]^, neural networks naturally lend themselves to analysis of single-cell datasets.

Single-cell RNA-Seq (scRNA-Seq) analysis follows similar strategies with bulk RNA-Seq where pre-processing is followed by normalization for library size and downstream analysis (see **Fig. 1** for a schema of a typical workflow). Due to the large number of cells being captured non-linear dimension reduction techniques have been extensively used for clustering and trajectory identification such as t-SNE and UMAP^[27,28]^. In addition to dimension reduction methods, scRNA-Seq datasets can be zero-inflated due to increased technical noise, hence various imputations approaches have been proposed. Furthermore, a general trend in scRNA-Seq analysis is to filter out genes that show relatively low variation across the dataset and filter out cells that express low number of genes. Although this is a valid strategy similar to bulk RNA-Seq analysis, the cutoff for number of top varying genes to select and number of genes expressed are generally arbitrary chosen, hence we aim to evaluate the effects of filtering genes and cells based on different thresholds as well. For this purpose, we combine various methods for different levels of analysis in a combinatorial fashion and evaluate identified trajectories in terms of cell orderings. Furthermore, since the ground-truth trajectories do not necessarily associate linearly with time in heterogeneous processes (e.g drug resistance), we have profiled clustering performance as well^[21]^.

**Figure 1.**
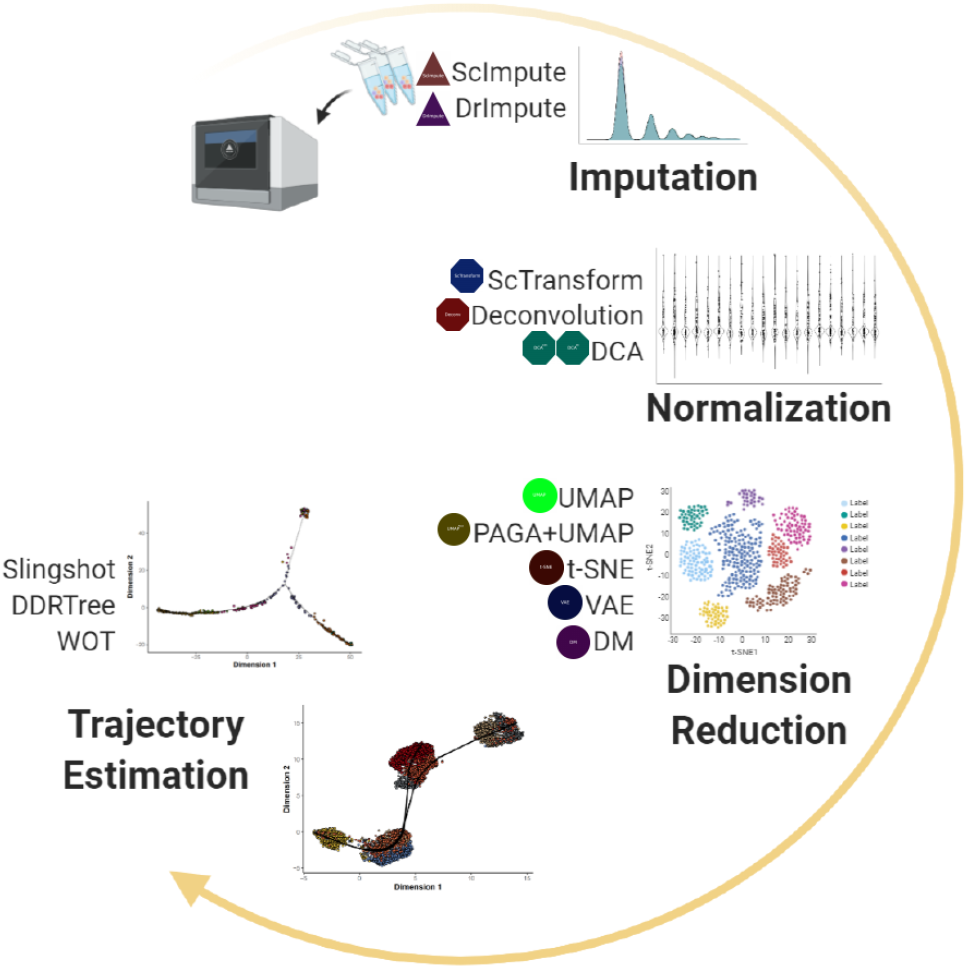
Schematic of general analysis steps and methods used for combinatorial workflows. Quality filtered raw read counts are processed through a step to reduce possible zero-count inflation by **one of** 2 **imputation methods**; ScImpute, DrImpute (or no imputation). Preprocessed data is **normalized by** 3 **methods**; ScTransform, Deconvolution and DCA followed by **dimension reduction using** 5 **methods**; UMAP, UMAP+PAGA, t-SNE, VAE, DM. Finally, **one of** 3 **trajectory inference methods** is used; Slingshot, DDRtree and WOT. Overall we have utilized 5376 analysis including the data subsets. (Note that the icons representative of individual methods are used to ease the interpretability of combinatorial workflows in downstream figures.)

We have utilized both publicly available and previously generated in-house dataset with variable number of cells, depth and complexity of the underlying process (**Table**.1). **TKI Treatment** dataset was previously generated to investigate transcriptional dynamics of resistance evolution to 3 TKIs; Alectinib, Lorlatinib and Crizotinib in lung cancer. In order to generate scRNA-Seq data with temporal information, cells were sampled at 4h (Alectinib only), 48h, 3w and 20-24w and sequenced. As we have hypothesized, this dataset represents a biologically heterogeneous example of evolutionary process hence crucial to evaluate PTEs. The **Pancreatic cell maturation** dataset contains transcriptional profiles of *α* and *β* cells during maturation process at 7 time-points; embryonic day 17.5 and postnatal days 0, 3, 9, 15, 18, and 60 representing relatively more homogeneous differentiation process. **Neurodegeneration** dataset is generated to investigate transcriptional dynamics of microglial cells isolated from Hippocampus at weeks 0, 1, 2, and 6 in CK-p25 inducible mouse model. **E2 Treatment** temporal scRNA-Seq is performed on 2 cell lines (MCF7,T47D) during 17*β* -estradiol (E2) treatment at 0h, 3h, 6h, and 12h.

**Table 1.** Datasets utilized in the study where the number of cells and genes are given prior to subset generation after quality control

| Dataset | Subset | Size (cells/genes) | Platform |
| --- | --- | --- | --- |
| **TKI Treatment**<sup>[39]</sup> | Alectinib | 5000/14000 | 10x |
|  | Lorlatinib | 4000/14000 | 10x |
|  | Crizotinib | 3700/14000 | 10x |
| **Pancreatic Maturation**<sup>[40]</sup> | $\alpha$ cells | 250/21700 | SmartSeq2 |
| | $\beta$ cells | 410/20500 | SmartSeq2 |
| **E2 Treatment**<sup>[41]</sup> | MCF7 | 60/21395 | Fluidigm C1 |
|  | T47D | 60/21570 | Fluidigm C1 |
| **Neurodegeneration**<sup>[42]</sup> | – | 800/15545 | SmartSeq2 |

Each dataset is preprocessed with different gene- and cell-level quality thresholds to generate 12 subsets and the overlap in estimated trajectories is quantified using rank correlation. For comparison of the effect of imputation we have used ScImpute, DrImpute which showed improved performance in various datasets and Deep Count Autoencoder (DCA) an autoencoder model aiming to combine de-noising and imputation in a single step^[15,16,29]^. We use **2 methods for normalization**; Deconvolution and ScTransform^[30,31]^. For DCA, since gene-wise dispersion and mean parameters are already estimated, we only used library size normalization. As scRNA-Seq clustering is an important step utilized in analysis of various datasets, we wanted to evaluate how robust are the clustering results when different workflows are used as well. For this purpose we coupled the Leiden clustering with **5 dimension reduction techniques**; UMAP, PAGA+UMAP, t-SNE, VAE and Diffusion Maps (DM) and evaluated the overlap of clusters using adjusted rand index (ARI)^[27,28,32,33,34,35]^. We have utilized **3 different trajectory inference methods**; Slingshot, DDRTree and WOT^[36,37,38]^. However, Slingshot operates on dimension reduced space hence we combined different dimension reduction methods with Slingshot as well. DDRTree on the other hand generates cell orderings by reducing the high-dimensional data to low-dimensional prinicple-tree structure, hence we have coupled DDRTree with preprocessing and normalization steps only^[37]^. Furthermore, since Slingshot and DDRTree are unsupervised approaches, we have utilized Waddington-OT (WOT), a supervised approach which aims to find cell-cell transition probabilities at consecutive time-points via optimization of unbalanced transcriptional mass transfer^[38]^. Comparison is somewhat imperfect however, as trajectories are defined slightly different for each method. Since slingshot estimates the smooth principle curve in low dimensional space, mapping of individual cells on the curve readily defines an ordering via the arc-length along the curve. In contrast, DDRTree embeds high-dimensional transcriptomic profiles onto a principle tree structure where the ordering is defined by the geodesic distances between individual cells. The supervised approach, WOT, on the other hand, generates a probability distribution between an individual cell at time *t*_*i*_ and the cell population at time *t*_*i*+1_, hence the trajectory of an individual cell is defined as the vector of transition probabilities. In order to evaluate the results from different methods in a comparable fashion, we opted to use Spearman’s rank correlation which does not take into account the distances between individual cells, but rather only the orderings, hence different quantitative scales between methods can be compared.

## Results

### Dimension reduction & Clustering

In order to evaluate how dimension reduction methods perform when coupled with Leiden method for clustering we have compared the identified clusters using Adjusted Rand Index (ARI) across different subsets of gene and cell level thresholded datasets. However, note that since we do not have ground-truth observations of clusters, instead we have focused on the overlap of identified clusters between different methods to assess repeatability. As expected we observed positive correlation of ARI across different workflows with the number of cells (**Fig**. S2). ARI values showed reduced overlap however between different methods across datasets globally, even when the number of cells is high (*ARI* < 0.75), suggesting that the selection of pre-processing methods for clustering purposes is non-trivial and can result in different cluster assignments. Investigating methods individually showed t-SNE as a method of choice where use of different preprocessing steps did not reduce overlap in the TKI treatment dataset and showed variable performance in remaining datasets in contrast with other methods (**Fig**. S3). Interestingly however neural network based methods showed variable results where use of DCA(NB/ZINB) as a preprocessing step in TKI dataset led to improved overlap between UMAP, UMAP+PAGA and t-SNE. In contrast, use of VAE as a dimension reduction method showed poor performance resulting in variable cluster assignments (**Fig** S4). This suggests that as a dimension reduction method, neural network based methods might not be the optimal choice but as a preprocessing step neural networks can provide advantages given that the number of cells is relatively high. Datasets with relatively low number of cells showed variable results. **F**or instance, in **P**ancreatic islet *α* cell differentiation, **UMAP, PAGA+UMAP and t-SNE showed improved overlap when D**rImpute is combined with **D**econvolution but overlap is reduced when Sc**T**ransform is used for normalization. **T**he neurodegeneration dataset on the other hand benefited from **DCA** with or without zero inflation model but overall showed decreased overlap (**Fig**. S3). These observations suggest that there is no one-size-fits-all solution for clustering of scRNA-Seq datasets and selection of parameters for preprocessing; normalization and dimension reduction can effect the clustering results hence methods required for analysis should be carefully selected especially when the number of cells is relatively low. If the number of cells is relatively high however, use of either DCA as a preprocessing step and/or t-SNE as a dimension reduction step improves overlap of clusters identified with different workflows.

### Trajectory Estimation

In order to evaluate PTEs mapping to a latent biological process we used Spearman rank correlation and normalized entropy. As given previously, using rank correlation we aim to do comparison of cell orderings identified by different workflows and normalized entropy is used to assess the distribution of rank correlations (bimodal around 0-1 or unimodal) in the case of Slingshot since > 1 PTEs are identified.

#### Slingshot

Evaluating the trajectories identified by Slingshot, we have observed large variation across different workflows and across different subsets. For instance, in Pancreatic maturation datasets, workflows that show relatively good overlap in *α* cell trajectories failed to identify overlapping trajectories in *β* cell dataset. Specifically, use of DrImpute or ScImpute resulted in decrease overlap of PTEs in *β* cell dataset (**Fig**. 2). Furthermore number of cells did not correlate positively with the repeatability of identified trajectories where majority of the workflows showed high entropy of rank correlations in TKI treatment dataset with minimum entropy being > 0.7 across 3 treatments (**Fig**. S5a,S5b,S5c). In contrast, datasets with relatively low number of cells showed slightly improved overlap for specific workflows. For instance in E2 treatment dataset use of DM improved overlap in contrast with UMAP or UMAP+PAGA. The neurodegeneration dataset on the other hand showed global decrease in PTEs (**Fig**. S5).

These results point out one of the major drawbacks of using Slingshot for PTEs; since estimation of trajectories is heavily dependent on the prior dimension reduction step, heterogeneous datasets will necessarily show high variation to different parameter regimes. More specifically, the Slingshot method using principle curves can fail to capture the temporal dynamics on highly non-linear spaces hence need to be carefully selected/optimized for trajectory estimation. For instance when UMAP is used for dimension reduction, the cell population structures remain overall similar as the number of cells increase but relative positioning of subpopulations can change in a way that does not reflect the latent temporal process (**Fig**. S6a,S6b,S6c,S6d). Furthermore the non-linearity can artificially generate increased number of trajectories leading to diverge PTEs. For instance, use of DM resulted in 1 trajectory to be identified in E2 treatment data subsets hence resulting in ‘simpler’ PTEs overall (**Fig**. S6g,S6h). This drawback was also noted in the original publication where Slingshot is suggested to be most appropriate when the temporal process is relatively linear and/or when there is a single trajectory. In order to alleviate some of the issues in coupling dimension reduction methods with Slingshot, one may need to choose parameter regimes towards linearity, for instance increasing number of nearest neighbors or minimum distance parameter in UMAP. This further stresses the importance of utilizing data specific methods and optimized parameters.

**Figure 2.**
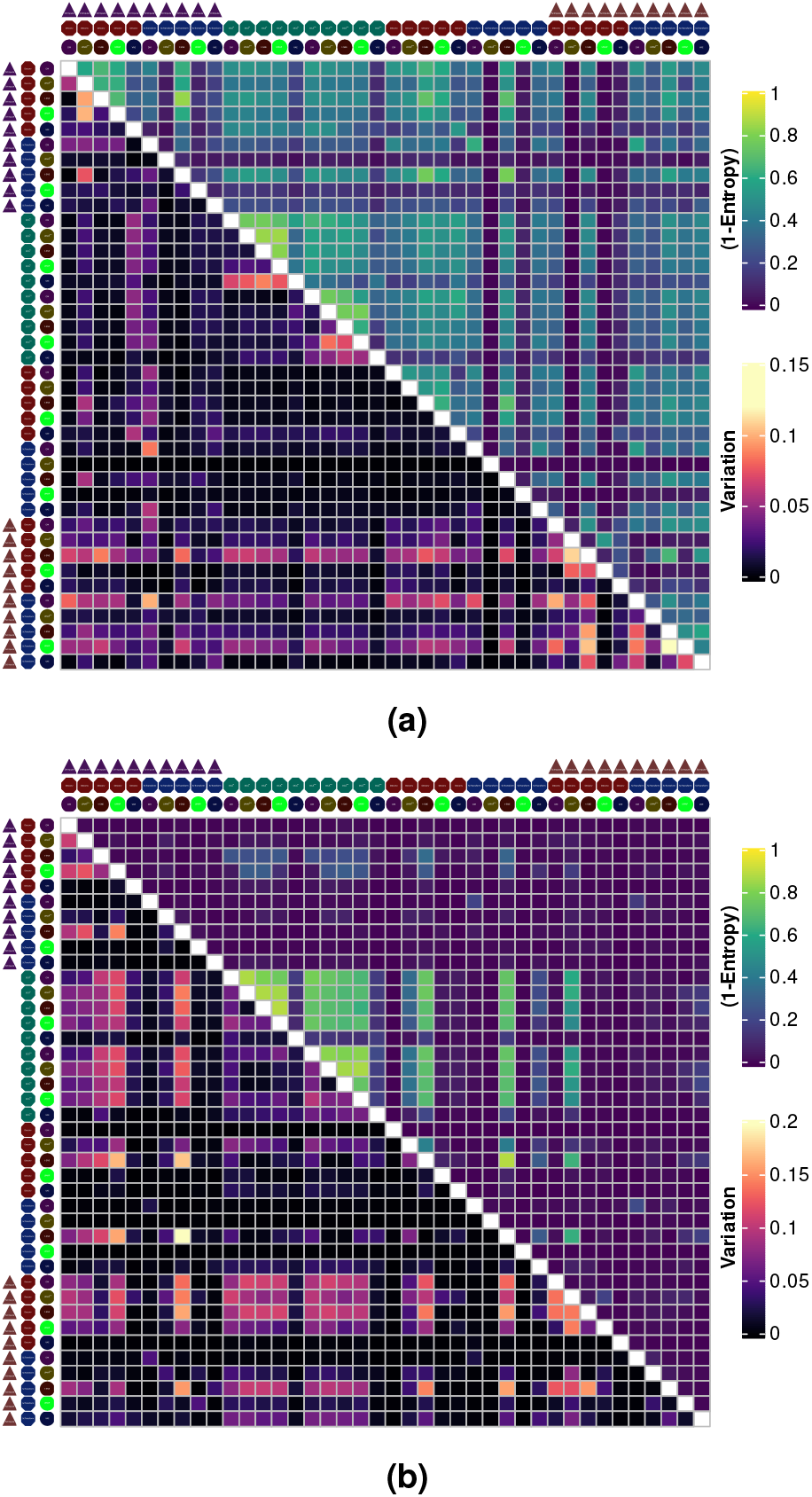
Comparison of trajectories identified by Slingshot showing data dependent performance of each workflow. 1-Entropy is used to aggregate the overlap between 2 trajectory quantified by rank correlation (upper-triangle). Better overlap results in lower entropy (See Supplementary for details). (a) Pancreatic maturation *α* cells, (b) Pancreatic maturation *β* cells.

#### DDRTree

Since DDRTree/Monocle2 method inherently utilizes dimension reduction to generate a tree-like topology to define a latent trajectory, we have generated the combinatorial workflows for imputation and normalization steps only. We have aggregated rank correlations across 12 subsets by median values to evaluate the overlap of different workflows. (**Fig**. S7). The TKI treatment dataset overall showed good overlap (*ρ* > 0.75) across different imputation and normalization methods. Interestingly however, Crizotinib treatment showed decreased overlap of PTEs when DrImpute or ScImpute is utilized in comparison with when DCA is used (**Fig**. 3a,3b,3c). Specifically, PTEs show increased correlation when DrImpute or ScImpute is used in contrast with when DCA is used. Further investigating the individual trajectories showed that using DCA resulted in increased number of branch points in contrast with DrImpute or ScImpute (**Fig**. S9). This might be an implication for ‘overcorrection’ when DrImpute or ScImpute is used subsequently reducing variation. Datasets with relatively low numbers of cells however showed variable results with different analysis steps having distinct ‘importance’. For instance, in the Neurodegeneration dataset, choice of normalization showed the highest impact where use of Deconvolution decreased the trajectory overlap globally, in contrast ScTransform was more robust to prior imputation step (**Fig**. 3d). Furthermore as expected, E2 treatment dataset showed high correlation between workflows using Deconvolution and ScTransform normalization but not when DCA is used (**Fig**. 3h, 3g) since with low number of training dataset for the autoencoder model, parameters might not be estimated robustly. In contrast, pooling information from similar cells and genes might better capture biological signal. The Pancreatic differentiation dataset on the other hand showed increased overlap across different methods. Further investigating subset specific overlap of trajectories showed no substantial effect of gene or cell level quality filtering where the quality of overlap between different workflows remained similar across different subsets (**Fig**. S8).

**Figure 3.**
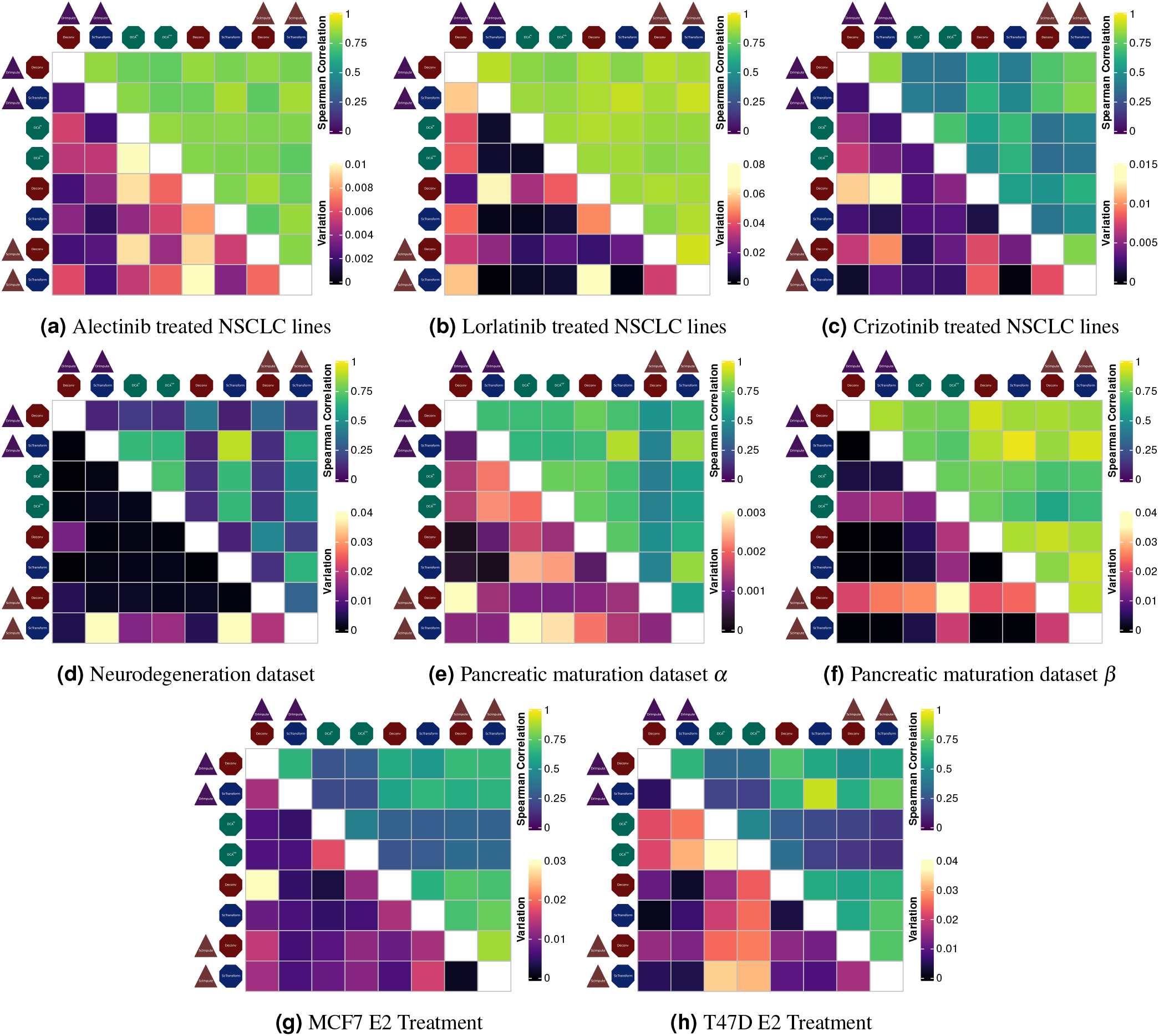
Rank correlation of geodesic distances on DDRTree trajectories aggregated over subsets showing both data specific performance and overall increase based on number of cells. (a-c) shows TKI treatment dataset where majority of the workflows result in relatively high correlation. (d-h) Remaining datasets show variable results with Pancreatic maturation *β* performing comparable to TKI dataset.

#### WOT

Since both Slingshot and DDRTree aim to find low dimensional ordering of individual cells in an unsupervised fashion, temporal information is not readily utilized which can lead to biased estimates where transcriptional dynamics are not ‘linearly’ associated with time (**Fig. 4**). Instead, supervised approaches can provide certain advantages for PTE by utilizing available temporal information hence capturing possible cyclic patterns where high transcriptional differences at initial time-points due to stress response for instance can diminish at later timepoints. However, forcing individual cells in a supervised order also poses challenges such that cells are not synchronized in terms of division and growth rates. For this purpose the WOT framework also allows us to calculate optimal growth rates for individual cells given the ‘transcriptional mass’ transfer optimization problem. Furthermore by removing the dimension reduction step, WOT inherently reduces the number of possible sources of variation. In order to evaluate how WOT performs when different methods for imputation and normalization are used, we have calculated pairwise rank correlations of transition probabilites between individual cells at consecutive time-points *t*_0_, *t*_1_, across different workflows. Simply, we have quantified how the transition probabilities of an individual cell change if a different normalization or imputation step is used. The TKI treatment dataset showed the highest overlap of transition probabilities across all pairwise workflow comparisons with median rank correlation > 0.75 (Fig. S10). Normalization with ScTransform showed slightly better overlap however when different imputation steps are compared in contrast with Deconvolution (**Fig**. S11). Interestingly however this difference was most striking in Neurodegeneration dataset where choice of imputation showed a relatively high difference of rank correlation (*ρ* > 0.2) between Deconvolution and ScTransform. Investigating imputation steps individually showed no substantial effect of imputation step where overlap of trajectories when Deconvolution and ScTransform is used remained similar and relatively low (< 0.75) irrespective of which imputation step is used (**Fig. S12**). Investigating the effect of using different gene and cell level thresholds showed a substantial decrease in the Neurodegeneration dataset where the remaining datasets showed similar PTE comparison profiles across 12 subsets hence suggesting relatively robust PTEs across different threshold (**Fig. S13**). This suggests that WOT PTEs consistently show repeatable results given that the dataset is large enough even for highly heterogeneous datasets.

**Figure 4.**
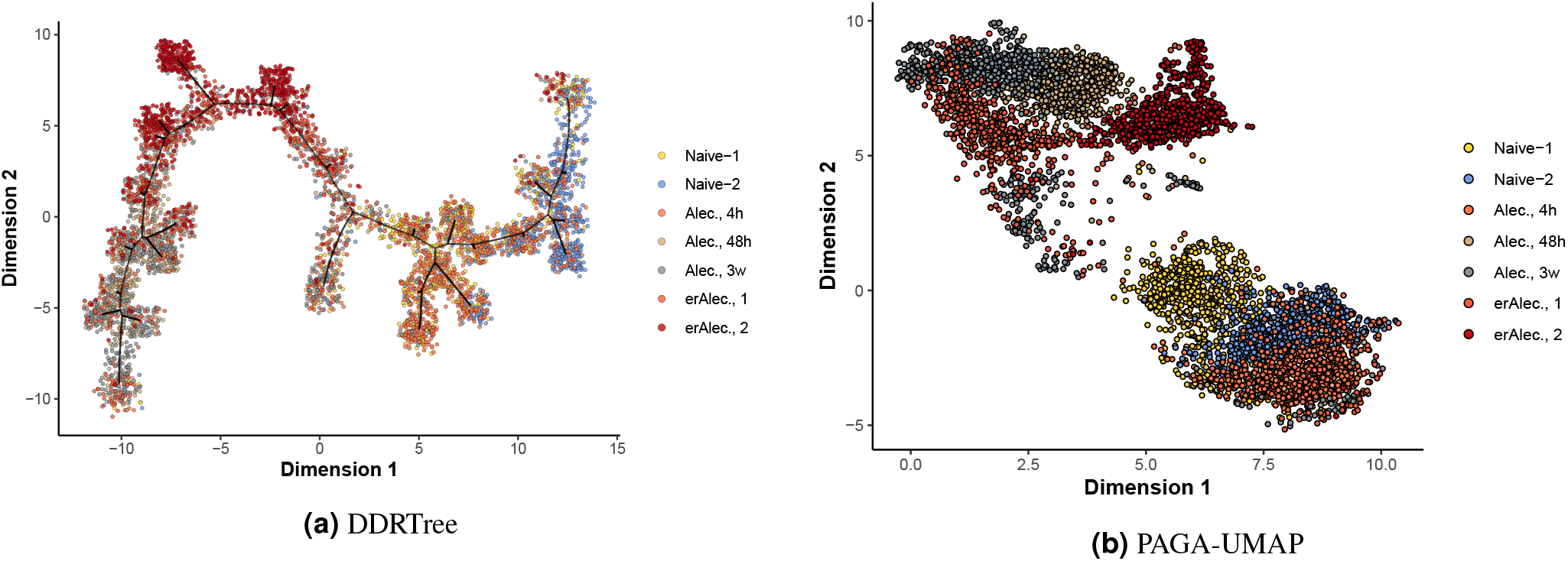
Sample dimension reductions for Alectinib treated NSCLC lines showing nonlinearity in temporal transcriptional dynamics. Since dimension reduction utilizes transcriptional similarity of individual cells, low dimensional representations might not necessarily correlate linearly with time where supervised approaches might be more suitable.

## Discussion

With the advent of single-cell sequencing methods, identification of tumor subpopulations and pseudotime estimation has been extensively used where analysis of scRNA-Seq data is complicated by a multitude of factors. In order to evaluate methods developed for scRNA-Seq analysis we have aimed at evaluating the available methods in a combinatorial fashion to assess the repeatability of either identified subpopulations or estimated pseudotimes. We have showed that selection of different methods at different levels of scRNA-Seq analysis can lead to variable outcomes both for clustering and trajectory inference. This is especially important considering the availability of additional methods not utilized in this study and the continued proliferation of methods. Furthermore, we have observed substantial variation in workflows for either clustering or PTE where non-linear dimension reduction methods are used. This emphasizes the importance of careful evaluation of which methods to utilize since the results may not be generalizable to replicate datasets.

General trends in our analysis showed that the number of captured cells is crucial when deciding on which downstream analysis methods to use since datasets with relatively high number of cells showed increased overlap across different workflows specifically for clustering and PTE using WOT. Dimension reduction methods that are heavily utilized in scRNA-Seq analysis showed high sensitivity to parameter selection hence clustering results using low dimensional representations were variable. Altough, t-SNE and UMAP coupled with PAGA showed better overlap there is no one best approach and methods showed data specific performance. This further stresses the importance of repeatability in scRNA-Seq analysis where hypothesis generation based on identified clusters embedded in low dimensional space is of major interest.

In order to alleviate some of the issues associated with clustering specifically when coupled with non-linear dimension reduction, ‘consensus’ based approaches where multiple clustering schemes are generated by randomly sampling features/cells might be more suitable. Similarly, trajectory inference methods showed variable results where non-linear dimension reduction is used. Slingshot for instance failed to capture reproducible trajectories in the TKI treatment dataset. As previously stated, Slingshot method based on principle curves is more suitable to relatively linear trajectories with small number of branch points. However, using either supervised method WOT or regularized dimension reduction using DDRTree resulted in increased correlations in trajectory estimates when different preprocessing methods are combined. DDRTree specifically showed improved performance over Slingshot especially when ScTransform is used for normalization but the quality of the overlap was data specific where TKI dataset with relatively large number of cells showed global increase in correlation of identified trajectories. Using supervised trajectory mapping via the WOT framework alleviated some of the issues with unsupervised approaches as well. Although identified trajectories remained sensitive to normalization method selection, data specific performance is reduced where we have observed ScTransform performing relatively well across all the datasets. Furthermore, since the temporal information is utilized in WOT, we can readily assume the identified trajectories will overlap with the biological process compared to unsupervised alternatives. For instance, neither Slingshot or DDRTree can differentiate subpopulations from different time-points if the transcriptional profiles are similar even though the temporal dynamics are different. However it is also important to note that identified trajectories only regard the differences between individual cells in terms of transcriptional profiles mapped to low dimensional space (in the case of Slingshot and DDRTree). This makes the problem of evaluating the PTEs non-trivial due to absence of ground-truth observations Consequently, as previously stated we have focused on the repeatability of PTEs. Deviation from ground-truth PTEs should be evaluated using approaches that allow individual cells to be tracked^[43,44]^. Furthermore, individual methods presented here can be further optimized separately resulting in global increase in PTE overlaps. For instance increasing the number of dimensions or using alternative metrics for quantifying transcriptional difference. Nevertheless, the WOT framework combined with ScTransform provided certain advantages by utilizing temporal information and reducing the variation.

In conclusion, analysis of scRNA-Seq datasets show high variation across different parameter regimes and methods in the context of clustering and trajectory mapping. It is non-trivial to utilize the heterogeneous structure of tumor subpopulations in order to extract biological insights hence analysis of scRNA-Seq requires careful selection of methods and optimization of parameters but different methods provide certain advantages. We hope that provided results can guide future studies for method selection and help with reproducibility in scRNA-Seq analysis.

## Appendices

### Methods

#### Imputation

##### ScImpute

ScImpute utilizes mixture modeling strategy to determine genes and cells that require imputation simultaneously. Using a Gamma-Normal mixture model, ScImpute first determines the dropout probability for each gene in each cell subpopulation identified by prior clustering. Imputation is then conducted using non-negative least squares regression using expression values from similar cells. We have used default parameters with dropout threshold set to 0.5 and initial clustering is done using *quickCluster* function from *scran* package in R with clustering method set to *igraph* and minimum size set to %10 number of cells.

##### DrImpute

DrImpute generates imputed counts using clustering and expression averaging where the cluster configurations are bootstrapped to result in robust estimates. All the parameters are set to default values.

#### Normalization

##### ScTransform

ScTransform utilizes a modified negative binomial (NB) regression where regularization using kernel-smoothing across mean expression levels for NB parameters is used. Passing library size as a covariate to NB regression allows for efficient normalization while accounting for mean-variance relationship. We have used *sctransform* package with parameter number of genes set to use all the genes.

##### Deconvolution

Normalization with deconvolution strategy aims to utilize count information from cells with similar transcriptional profiles. For this purpose, initially clustered cells are normalized against the population as a single pool of cells. The normalization factor is then deconvolved to generate cell-wise normalization factors using least squares methods. Similar to ScImpute, initial pool of cells are generated by using *quickCluster* method from *scran* package with clustering method set to *igraph* and minimum size set to %10

##### Deep Count Autoencoder

Deep Count Autoencoder (DCA) follows a slightly different strategy in which autoencoder based neural network is used to estimate dropout and dispersion parameters with likelihood based on negative-binomial (NB) or zero-inflated negative-binomial (ZINB) models are used. DCA aims to identify the latent structure in scRNA-Seq data leading to observed noise and dropouts by constraining the latent dimension to *d* << *p* where *p* is it total feature/gene space and *d* is the latent space. Since DCA estimates the mean expression parameter *µ* for each gene for each cell, we have used library size normalization on the *µ* parameters for both NB and ZINB models effectively generating 2 normalized datasets with imputed and non-imputed pre-processing.

#### Dimension Reduction

##### UMA

Uniform Manifold Approximation and Projection (UMAP) is a nonlinear dimension reduction technique heavily utilized in scRNA-Seq data analysis. For a detailed explanation see the original article and python package documentation ^[27,45]^. Simply UMAP aims to build a k-nearest neighbor graph defined by the parameter ‘number of neighbors’ and generates an ‘n’ dimensional representation where distances of k-nearest neighbors are preserved across the dataset. Where possible, we have used default parameters 30 for ‘n_neighbors’ and 0.5 for ‘min_dist’. For datasets with low number of cells we have set ‘n_neighbors’ as 10.

##### SNE

Similar to UMAP, t-Distributed Stochastic Neighbor Embedding (t-SNE) aims to find a low dimensional representation by minimizing KL-Divergence between pairwise distances in original space with low dimensional representation^[28]^. The perplexity parameter is set to 30 for datasets with high number of cells 10 otherwise. *θ* parameter is set to 0.01 as well.

##### UMAP+PAGA

partitioned-based Graph Abstraction (PAGA) generates a low dimensional k-nearest neighbor graph embedding utilizing cell-cell similarities. Using PAGA embedded coordinates, UMAP can then be initialized with the PAGA coordinates allowing to couple PAGA with UMAP aiming to increase representative power of reduced dimensions^[33]^.

##### DM

Diffusion Maps is a non-linear dimension reduction technique aiming to improve Principle Component based methods by incorporating ‘diffusion’ through construction of transition matrices. Eigenvectors of the constructed transition matrix generates the reduced subspace where *t* − *th* power will represent a random-walk like diffusion distance betwen individual cells^[35,46,47]^.

##### Variational Autoencoder

Variational Autoencoders (VAE) widely used as generative models where the encoder model embedds the parameters of data prior and the decoder model generates the data from points sampled from embedded distribution. More simply, the encoder network embedds the mean and dispersion parameters where the prior distribution is assumed to be isotropic gaussian with mean 0 and standard-deviation 1. Coupled with proper regularization, VAEs can effectively represent the given data in low dimensional space. The neural network is constructed to have 3 hidden layers with 1024, 512, 256 units for both the encoder and decoder network and a single stochastic latent layer with 2 units. We have used RMSprop optimizer with a learning rate of 0.0001

#### Trajectory Mapping

**Slingshot:** utilizes principle curves to generate cell orderings given the minimum. Briefly, Slingshot uses initial clustering of cells followed by minimum-spanning tree (MST) identification of the cell clusters. Identified MST structure generates a cluster lineage which is followed by principle curve fitting simultaneously for each branch in the MST^[36]^.

**DDRTree:** aims to find a regularized low dimensional projection (*d* << *D*) of original high-dimensional dataset onto a space formed by orthogonal set of basis vectors with minimum-spanning tree (MST) regularization term^[48,49]^. Also note that due to computational constraints, we have initially reduced the data to 50 principal components. Default values are used for remaining set of parameters.

**Waddington-OT:** is a supervised trajectory mapping method where sequencing time-points are used to construct transition probabilities between consecutive cell populations. Transition probabilities are estimated by optimizing an unbalanced optimal-transport problem where the aim is to find a mapping between cells minimizing the cost incurred by transcriptional mass difference (euclidean distance of transcriptional profiles)^[38]^. All the workflows and scripts used to run analysis are deposited online and publicly available at https://github.com/ardadurmaz/sc_eval.

##### Trajectory Comparison

We have extensively utilized rank correlation and where there are multiple trajectories entropy as a measure to quantify overlap of pseudo-time estimate (PTE) obtained from different workflows. Since Slingshot can generate > 1 trajectories in order to evaluate the distribution of spearman correlations between pairwise trajectory comparison, we calculated entropy to quantify whether spearman correlations are bimodal around 0-1 corresponding to ‘high’ quality overlap between trajectories (**Fig**.S1).

**Figure S1.**
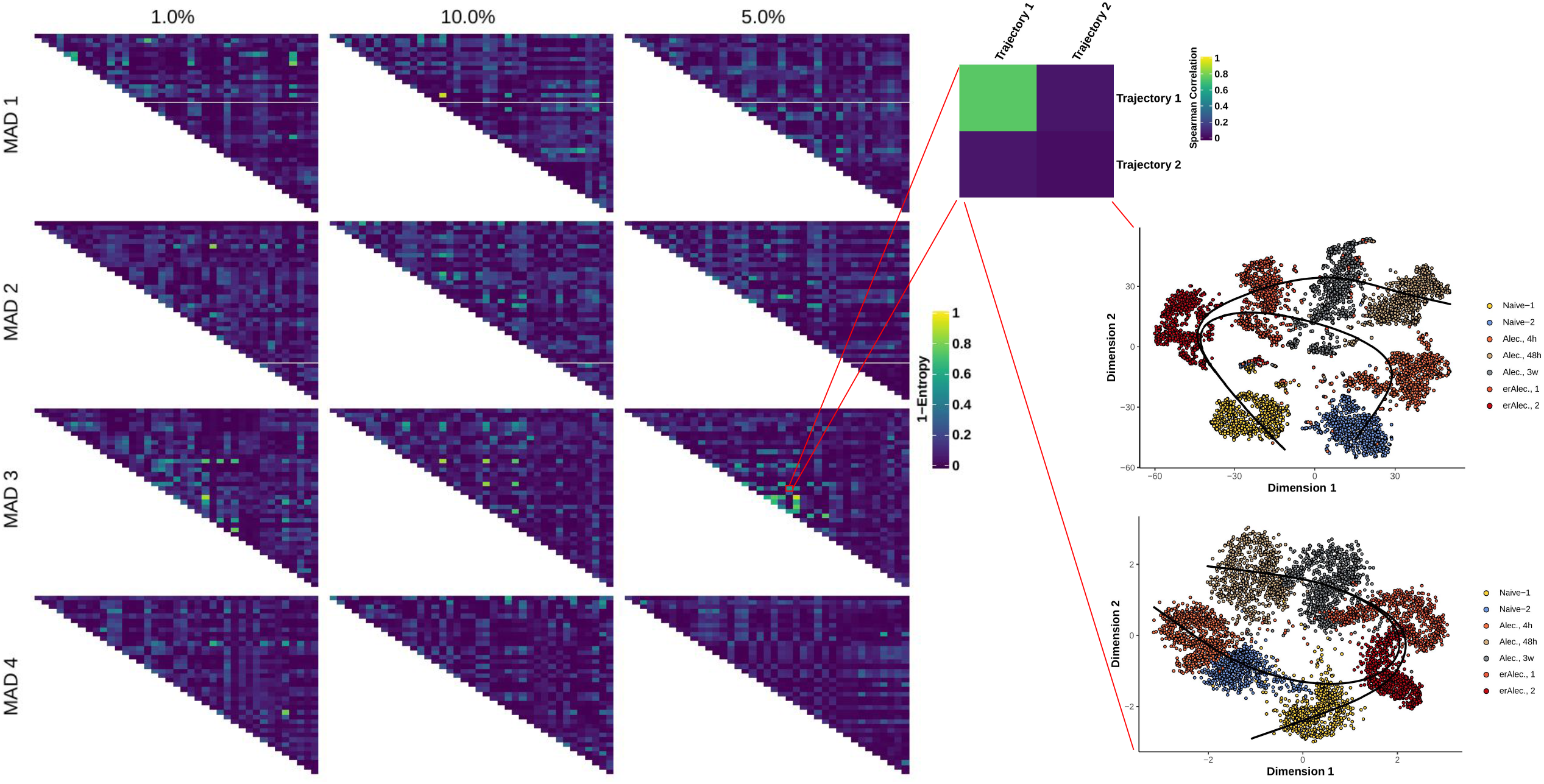
Example figure showing the quantification by entropy over multiple trajectories identified. In order to quantify the global overlap we have compared inidividual trajectories using spearman correlation scaled between 0-1. Using the scaled spearman correlations as pseudo-probabilities, we calculated entropy to assess whether scaled spearman correlations are centered around 0-1 suggesting good overlap.

**Figure S2.**
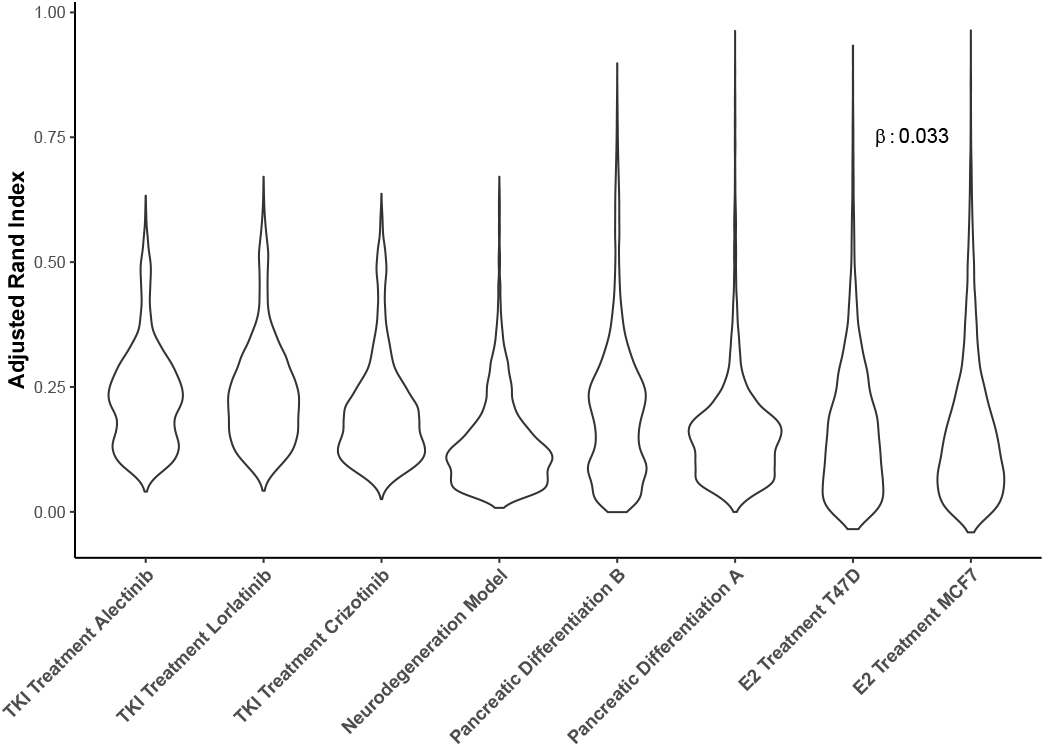
Adjusted rand index distribution across different datasets ordered by decreasing number of cells. Each point represents a pairwise comparison of clusters identified using different combinatorial workflows. Linear regression of ARI using number of cells as a covariate shows significant association with *β* = 0.03 (p< 0.001).

**Figure S3.**
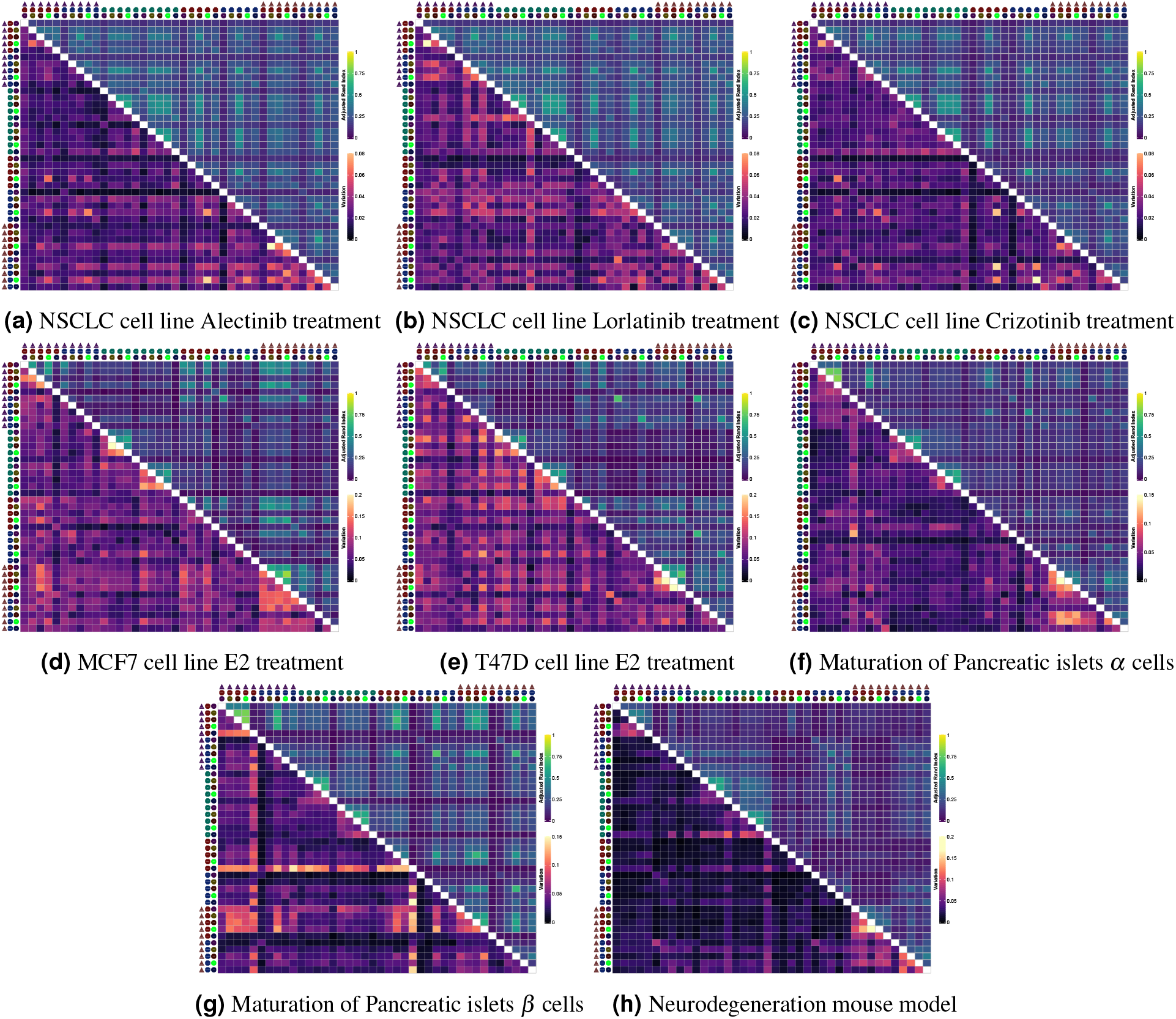
Comparison of clusters identified using Leiden clustering. Adjusted rank index across different methods is used to evaluate cluster overlaps which us further summarized by taking the median across 12 subsets.

**Figure S4.**
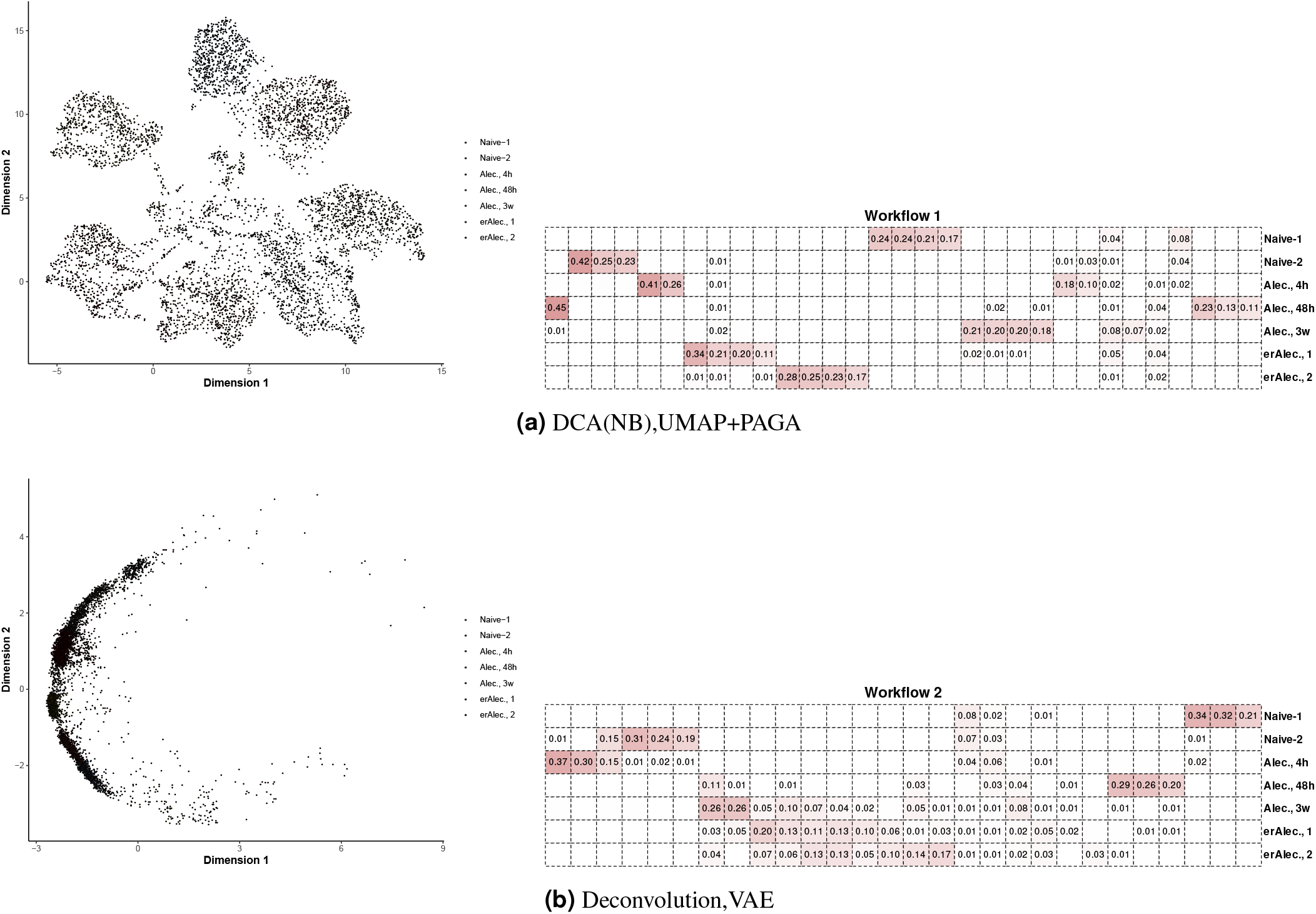
Sample cluster identification showing distinct results for both reduced dimensional representation and identified clusters between UMAP+PAGA and VAE dimension reduction methods. Heatmaps show the row-normaized percentage of cells at specific sampling time-point (rows) assigned to distinct clusters (columns) in 1 data subset out of 12.

**Figure S5.**
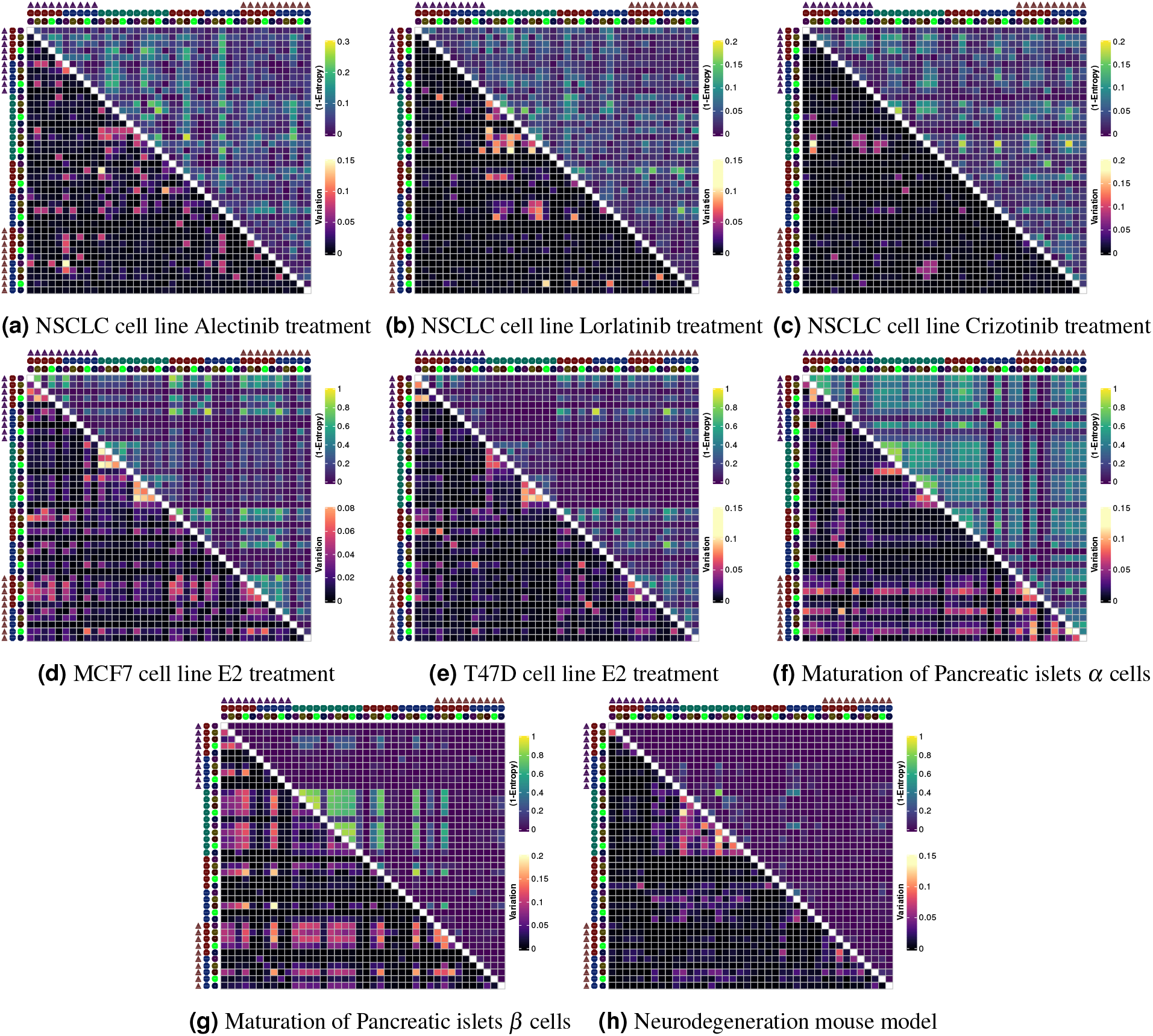
Comparison of trajectories identified by Slingshot. Quality of overlap is summarized by quantifying the ‘randomness’ of scaled Spearman rank coefficients between the trajectories. Treating scaled rank coefficients as pseudo-probabities and using entropy allowed us to assess whether the pairwise trajectory comparisons are bimodal around 0 and 1 (suggesting good mapping) or uniformly distributed (suggesting no optimal mapping). 1-Entropy values are then averaged across 12 subsets. Upper triangle shows the aggregated entropy values and lower triangle shows the variation in entropy values (Best overlap would be represented by low entropy and low variation values).

**Figure S6.**
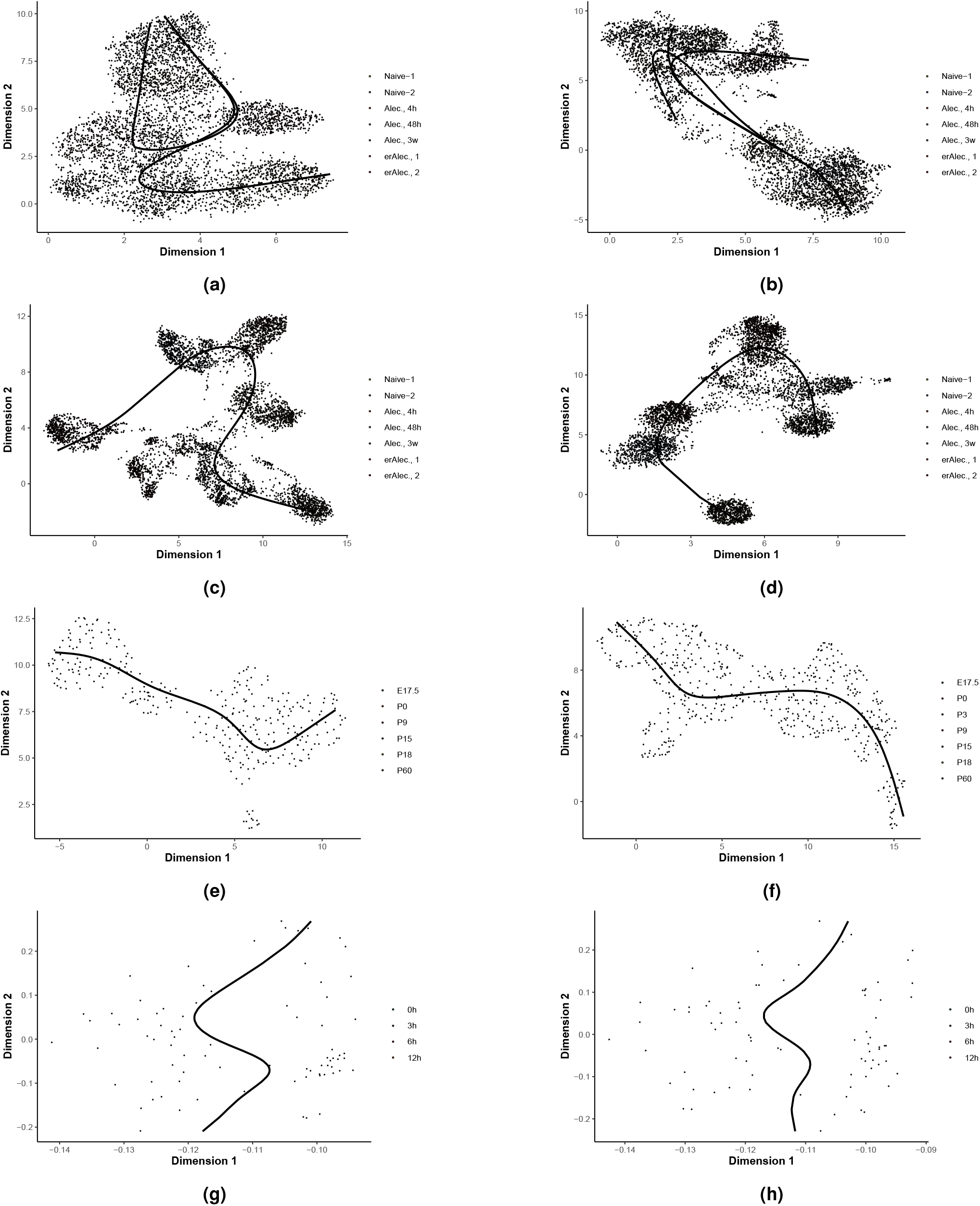
Example Slingshot trajectory estimates using (a-b) Deconvolution and ScTransform coupled with PAGA+UMAP on non-imputed dataset (c-d) Deconvolution and ScTransform coupled with PAGA+UMAP on imputed data with DrImpute. (e-f) shows Slingshot applied on Pancreatic maturation *α* and *β* cells respectively processed using DCA and dimension reduced with UMAP+PAGA showing relatively good overlap. (g-h) shows DM applied in E2 treatment dataset for DrImputed and no-imputation respectively.

**Figure S7.**
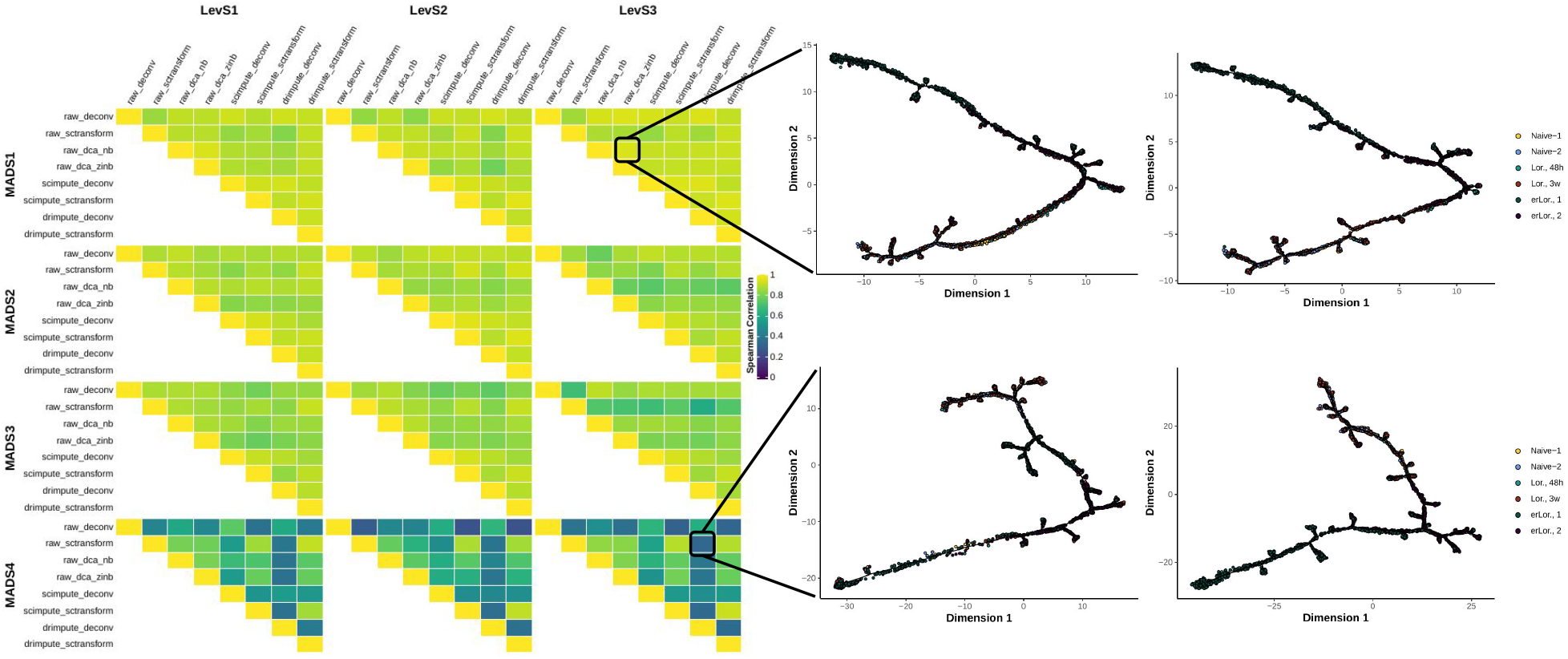
Comparison of pairwise trajectories for Lorlatinib treated NSCLC cell line separately for 12 subsets. Individual rank correlations are then aggregated by taking the median to summarize overall similarity of pairwise workflows.

**Figure S8.**
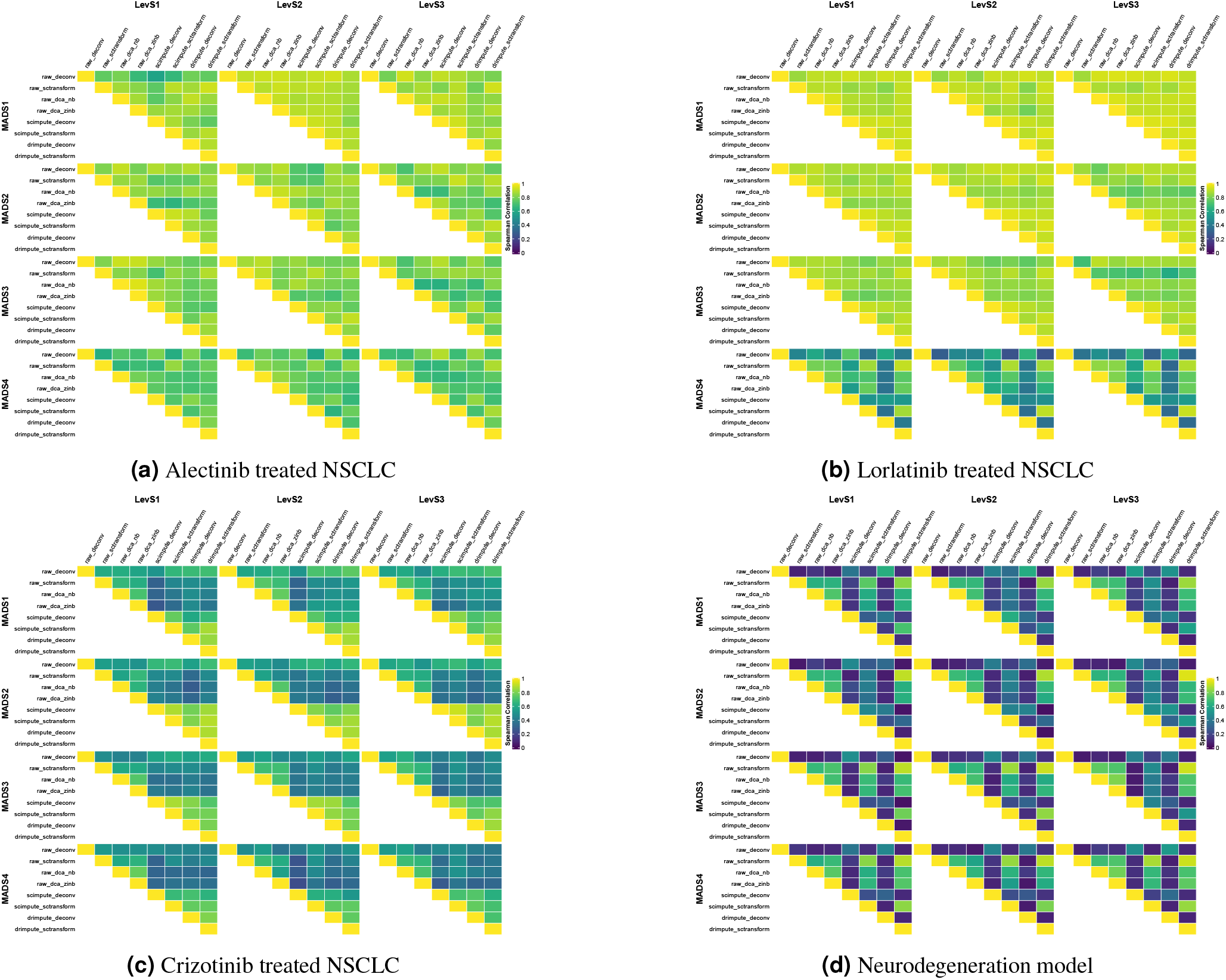
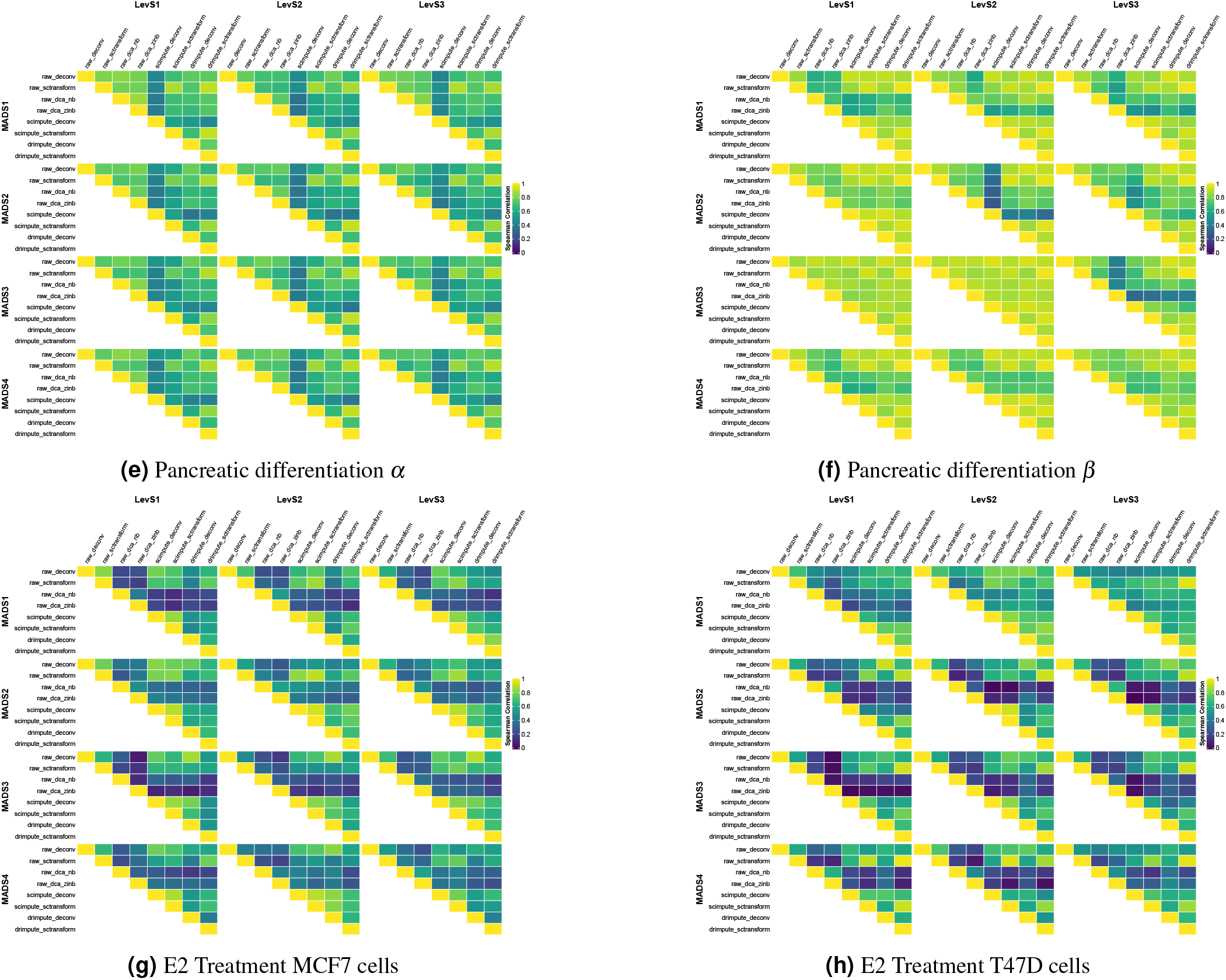
Overlap of DDRTree trajectories across different subsets stratified by cell level (X-Axis), gene level (Y-Axis) thresholds and workflows quantified by the spearman rank correlation of geodesic distances between individual cells. No substantial difference exists in rank correlations across different thresholds for pairwise workflow comparisons.

**Figure S9.**
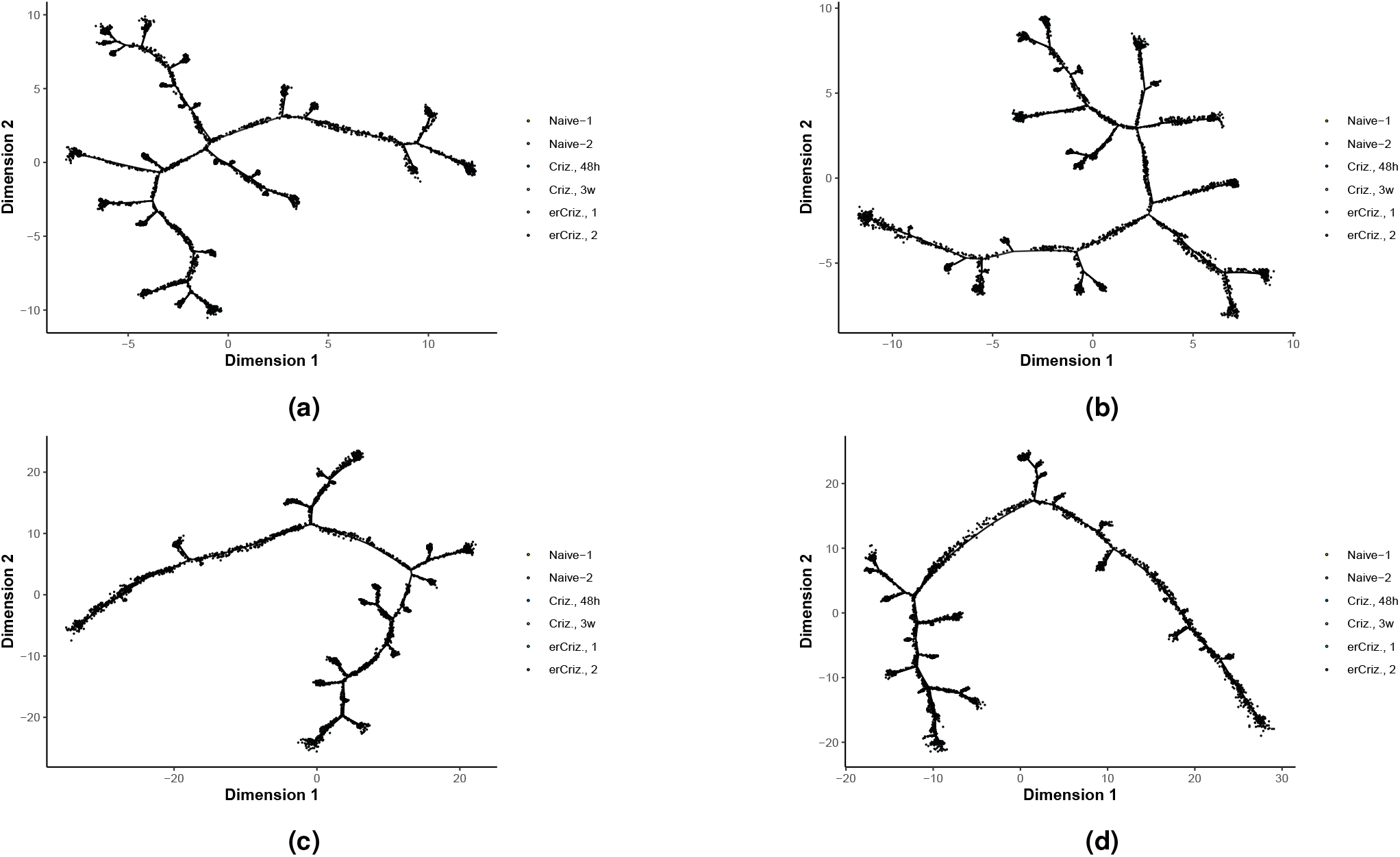
Trajectories identified by DDRTree using Crizotinib dataset showing increased number of branch-points identified when DCA is utilized. (a-b) shows DCA-NB and DCA-ZINB respectively, (c-d) shows applying DrImpute and ScImpute followed by ScTransform respectively

**Figure S10.**
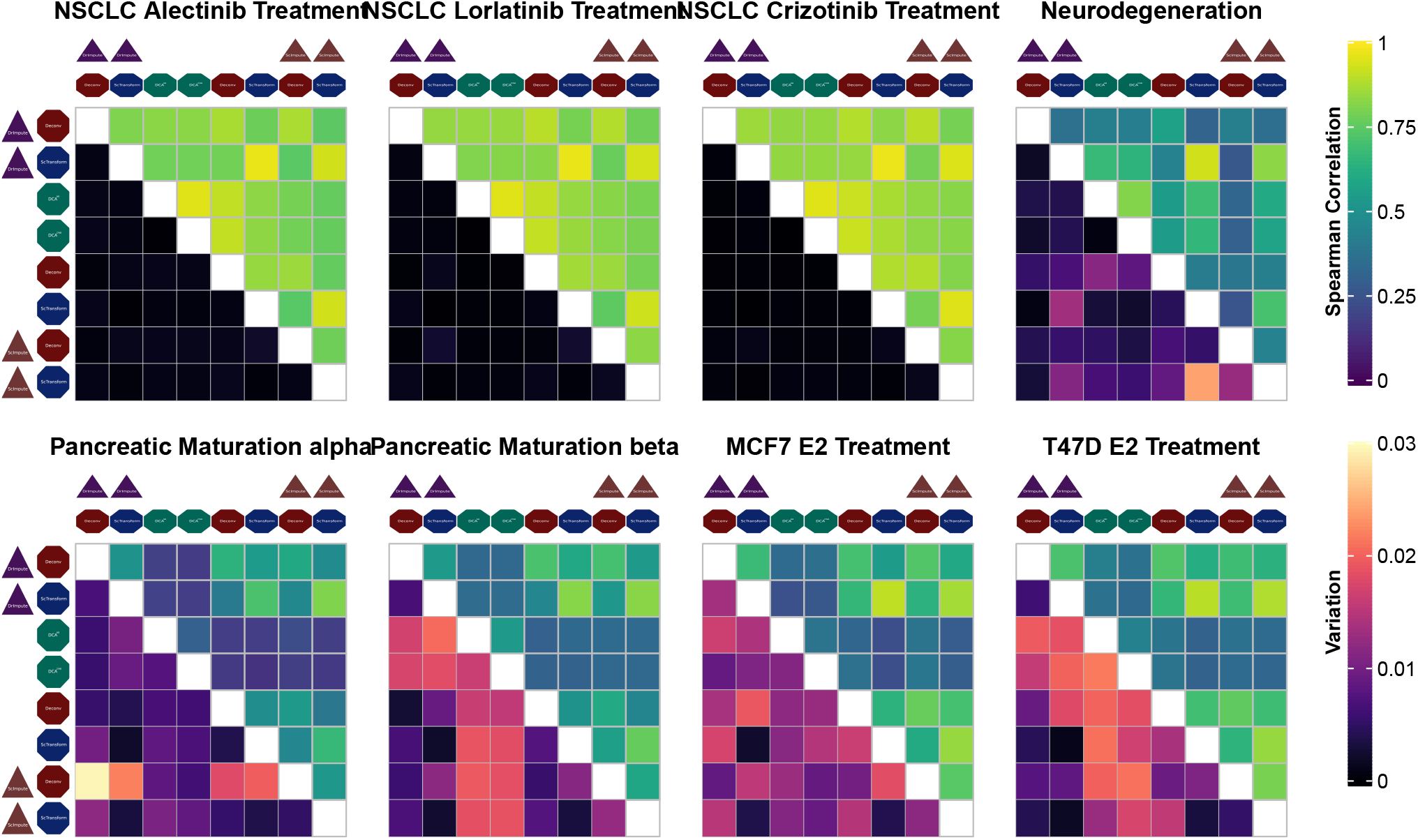
Waddington-OT PTEs comparisons showing median rank correlations across 12 subsets (upper-triangle) and associated variation (lower-triangle)

**Figure S11.**
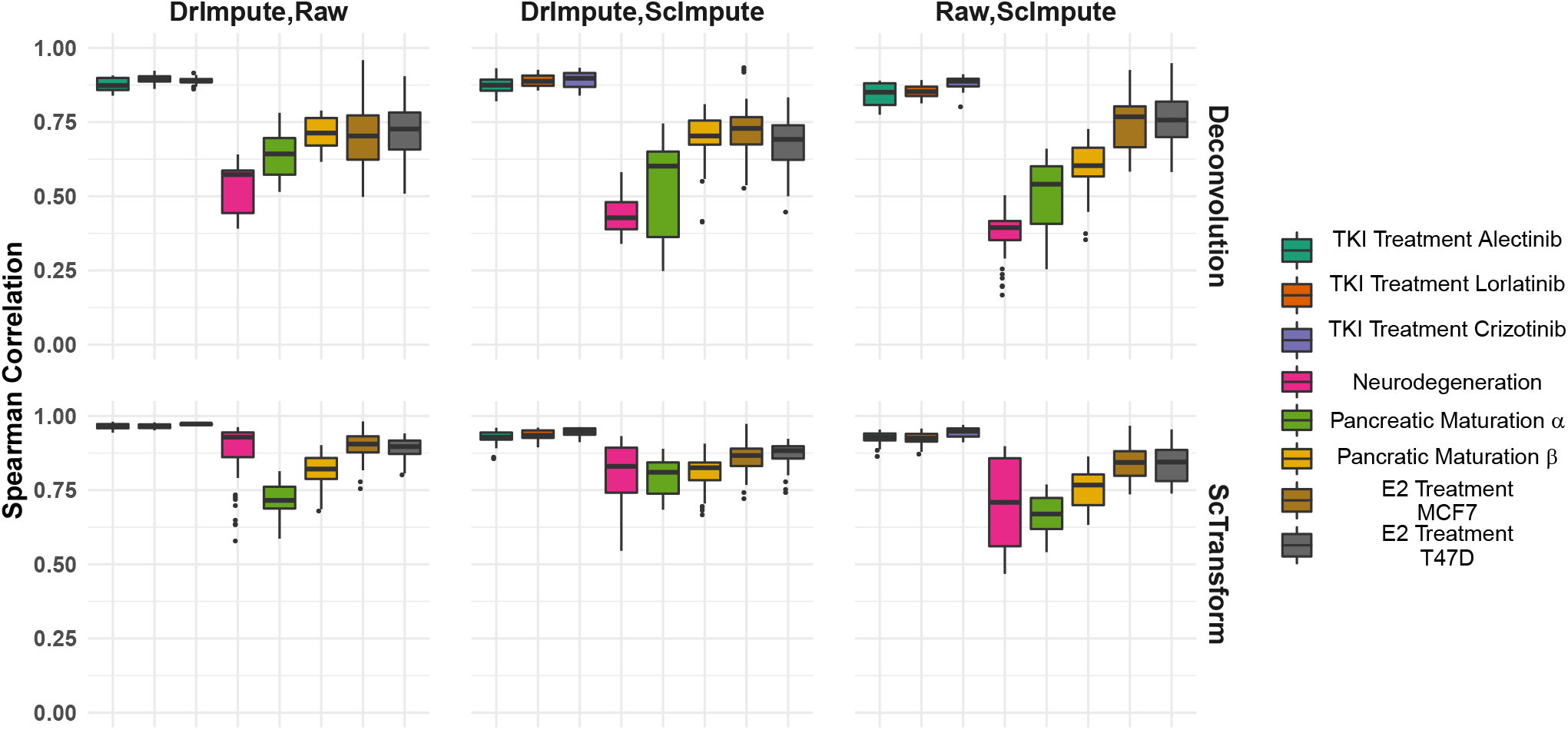
Waddington-OT rank correlation comparison for normalization methods ScTransform and Deconvolution showing a global trend towards improved ScTransform PTE overlaps. Individual points represent the rank correlation between different imputation workflows when Deconvolution and ScTransform is used as normalization step.

**Figure S12.**
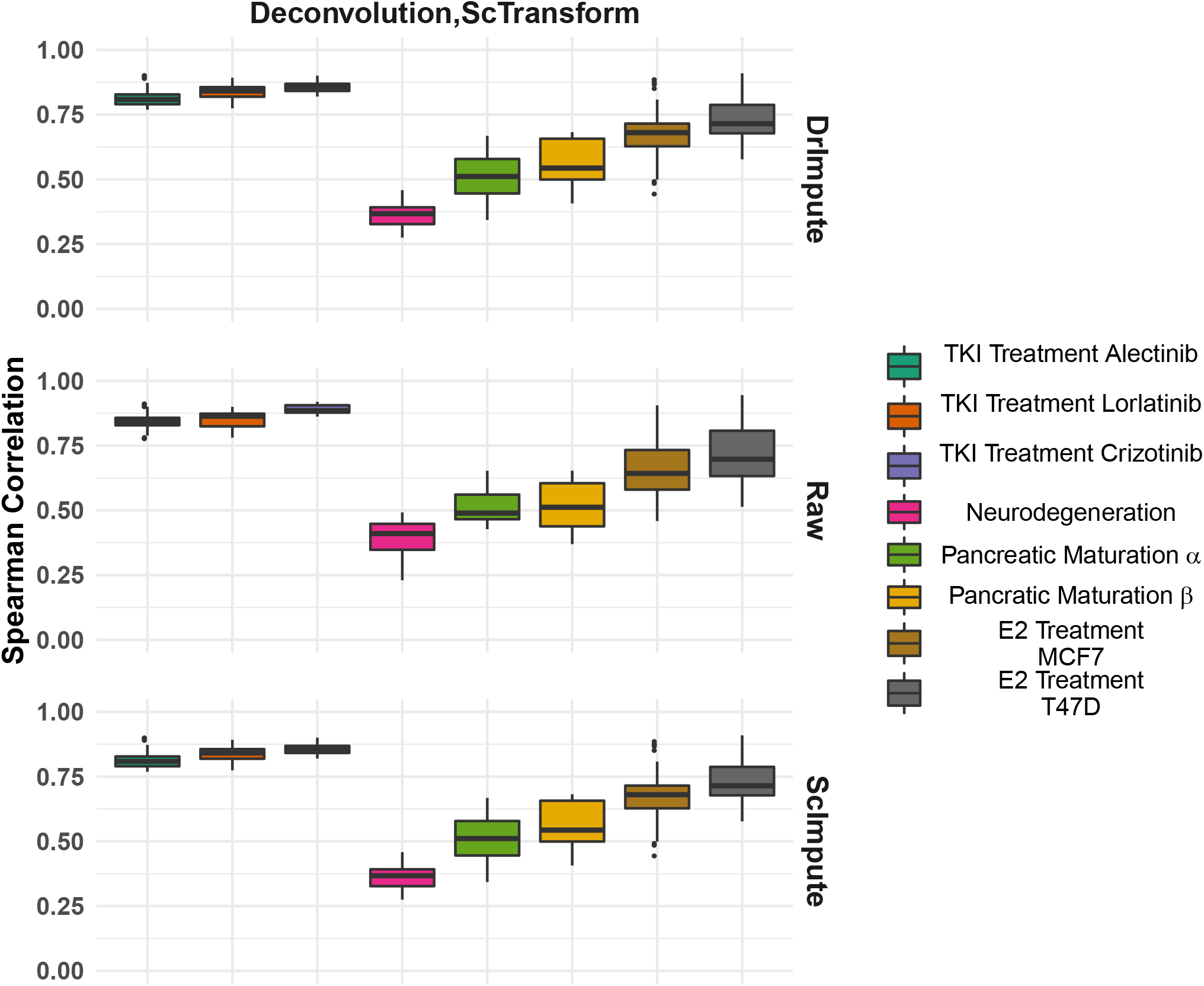
Comparison of Imputation methods showing no substantial effect of preprocessing on WOT PTEs. Using ScTransform and Deconvolution for normalization shows no substantial dependence on imputation step hence resulting in similar rank correlations

**Figure S13.**
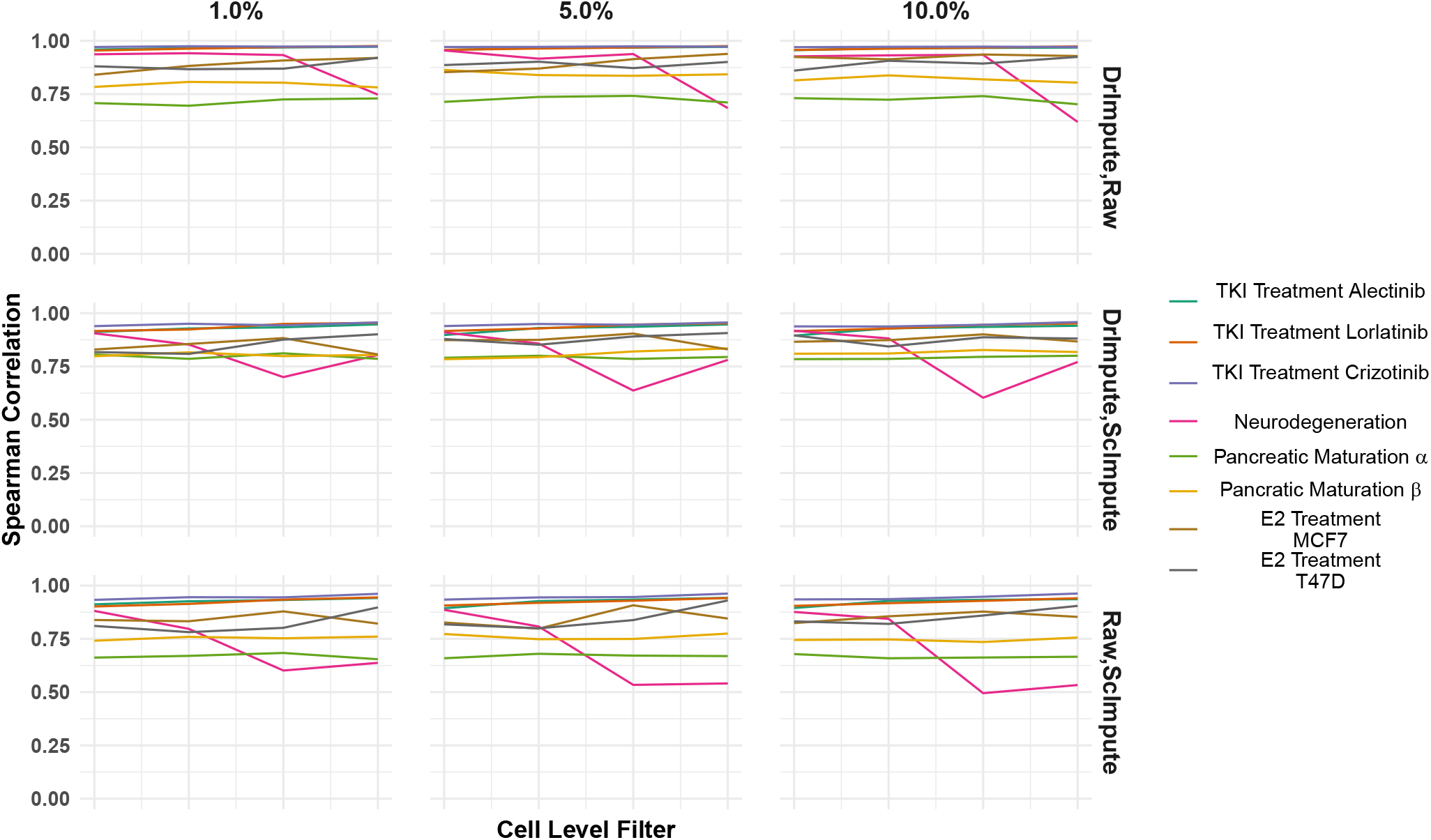
WOT PTEs comparisons when ScTransform is used for normalization across 12 subsets separated by cell level and gene level filtering showing reduced effect. Spearman rank correlations show no substantial difference when different thresholds are used for filtering out low quality cells and genes. x-axis is ordered in increasing cell level threshold and each facet is given in increasing order of gene level threshold (1%, 5%, 10%).

**Figure S14.**
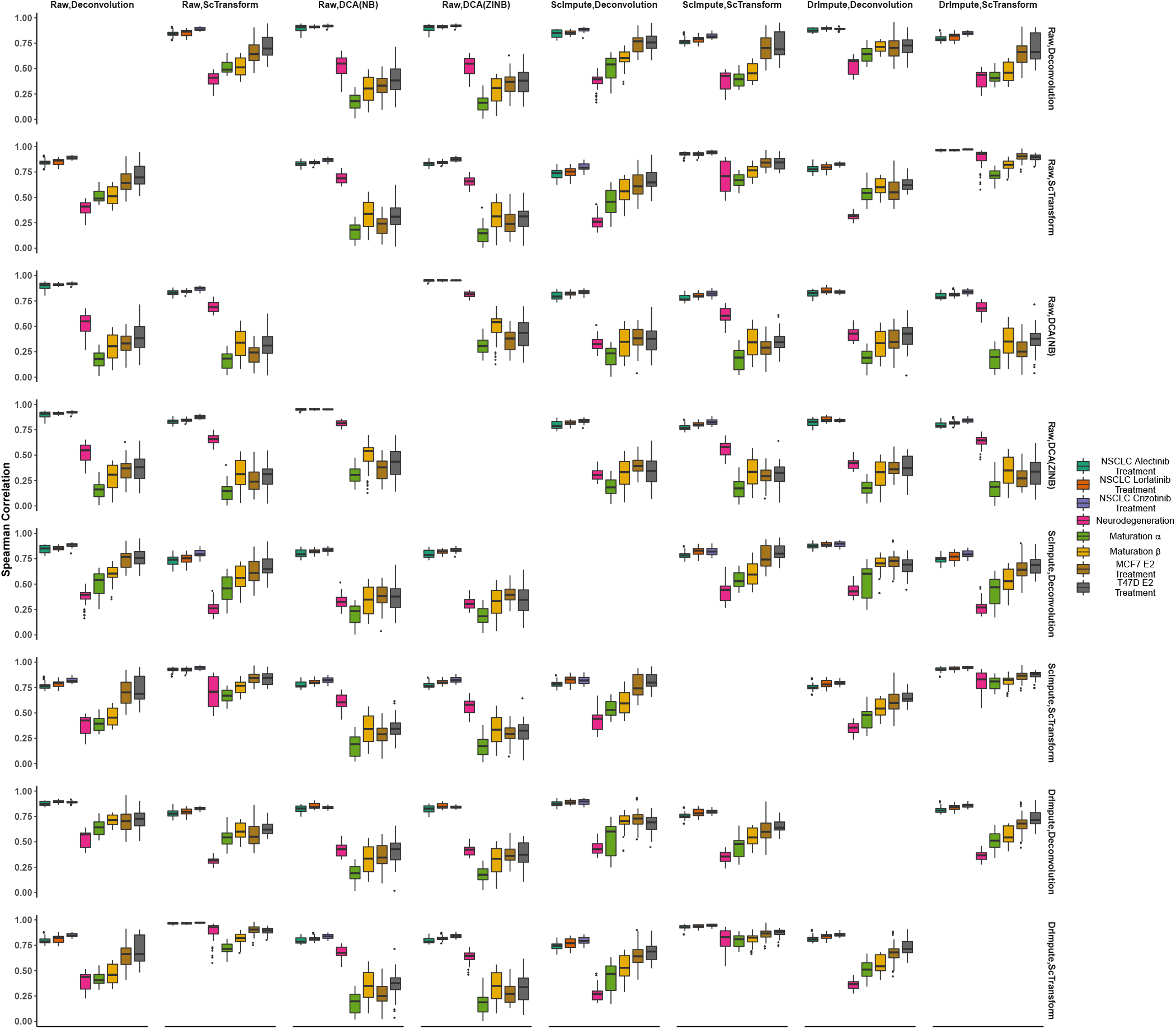
Distribution of Waddington-OT rank correlations between different workflows.

## References

1. Marusyk, A., Almendro, V. & Polyak, K. Intra-tumour heterogeneity: a looking glass for cancer? Nat. Rev. Cancer 12, 323 (2012).

2. Hinohara, K. & Polyak, K. Intratumoral heterogeneity: More than just mutations. Trends cell biology (2019).

3. Kreso, A. & Dick, J. E. Evolution of the cancer stem cell model. Cell stem cell 14, 275–291 (2014).

4. Burrell, R. A. & Swanton, C. Tumour heterogeneity and the evolution of polyclonal drug resistance. Mol. oncology 8, 1095–1111 (2014).

5. Quail, D. F. & Joyce, J. A. Microenvironmental regulation of tumor progression and metastasis. Nat. medicine 19, 1423 (2013).

6. Kaznatcheev, A., Peacock, J., Basanta, D., Marusyk, A. & Scott, J. G. Fibroblasts and alectinib switch the evolutionary games played by non-small cell lung cancer. Nat. ecology & evolution 3, 450–456 (2019).

7. Lee, M.-C. W. et al. Single-cell analyses of transcriptional heterogeneity during drug tolerance transition in cancer cells by rna sequencing. Proc. Natl. Acad. Sci. 111, E4726–E4735 (2014).

8. Kim, K.-T. et al. Single-cell mrna sequencing identifies subclonal heterogeneity in anti-cancer drug responses of lung adenocarcinoma cells. Genome biology 16, 127 (2015).

9. Tirosh, I. et al. Single-cell rna-seq supports a developmental hierarchy in human oligodendroglioma. Nature 539, 309 (2016).

10. Sharma, A. et al. Longitudinal single-cell rna sequencing of patient-derived primary cells reveals drug-induced infidelity in stem cell hierarchy. Nat. communications 9, 4931 (2018).

11. Hong, S. P. et al. Single-cell transcriptomics reveals multi-step adaptations to endocrine therapy. bioRxiv 485136 (2018).

12. Nichol, D. et al. Antibiotic collateral sensitivity is contingent on the repeatability of evolution. Nat. communications 10, 1–10 (2019).

13. Card, K. J., LaBar, T., Gomez, J. B. & Lenski, R. E. Historical contingency in the evolution of antibiotic resistance after decades of relaxed selection. PLoS biology 17, e3000397 (2019).

14. Lv, B. et al. Single-cell rna sequencing reveals regulatory mechanism for trophoblast cell-fate divergence in human peri-implantation conceptuses. PLoS biology 17, e3000187 (2019).

15. Li, W. V. & Li, J. J. An accurate and robust imputation method scimpute for single-cell rna-seq data. Nat. communications 9, 1–9 (2018).

16. Gong, W., Kwak, I.-Y., Pota, P., Koyano-Nakagawa, N. & Garry, D. J. Drimpute: imputing dropout events in single cell rna sequencing data. BMC bioinformatics 19, 1–10 (2018).

17. Saelens, W., Cannoodt, R., Todorov, H. & Saeys, Y. A comparison of single-cell trajectory inference methods. Nat. biotechnology 37, 547–554 (2019).

18. Tian, L. et al. Benchmarking single cell rna-sequencing analysis pipelines using mixture control experiments. Nat. methods 16, 479–487 (2019).

19. Yosef, N. & Regev, A. Impulse control: temporal dynamics in gene transcription. Cell 144, 886–896 (2011).

20. Steinacher, A., Bates, D. G., Akman, O. E. & Soyer, O. S. Nonlinear dynamics in gene regulation promote robustness and evolvability of gene expression levels. PloS one 11, e0153295 (2016).

21. Lee, M. J. et al. Sequential application of anticancer drugs enhances cell death by rewiring apoptotic signaling networks. Cell 149, 780–794 (2012).

22. Rumelhart, D. E., Hinton, G. E. & Williams, R. J. Learning internal representations by error propagation. Tech. Rep., California Univ San Diego La Jolla Inst for Cognitive Science (1985).

23. Kingma, D. P. & Welling, M. Auto-encoding variational bayes. arXiv preprint arXiv:1312.6114 (2013).

24. Palazzo, M., Beauseroy, P. & Yankilevich, P. A pan-cancer somatic mutation embedding using autoencoders. BMC bioinformatics 20, 1–10 (2019).

25. Xiao, Z. & Deng, Y. Graph embedding-based novel protein interaction prediction via higher-order graph convolutional network. PloS one 15, e0238915 (2020).

26. Ding, J. & Regev, A. Deep generative model embedding of single-cell rna-seq profiles on hyperspheres and hyperbolic spaces. BioRxiv 853457 (2019).

27. McInnes, L., Healy, J. & Melville, J. Umap: Uniform manifold approximation and projection for dimension reduction. arXiv preprint arXiv:1802.03426 (2018).

28. Maaten, L. v. d. & Hinton, G. Visualizing data using t-sne. J. machine learning research 9, 2579–2605 (2008).

29. Eraslan, G., Simon, L. M., Mircea, M., Mueller, N. S. & Theis, F. J. Single-cell rna-seq denoising using a deep count autoencoder. Nat. communications 10, 1–14 (2019).

30. Lun, A. T., Bach, K. & Marioni, J. C. Pooling across cells to normalize single-cell rna sequencing data with many zero counts. Genome biology 17, 75 (2016).

31. Hafemeister, C. & Satija, R. Normalization and variance stabilization of single-cell rna-seq data using regularized negative binomial regression. Genome Biol. 20, 1–15 (2019).

32. Traag, V. A., Waltman, L. & van Eck, N. J. From louvain to leiden: guaranteeing well-connected communities. Sci. reports 9, 1–12 (2019).

33. Wolf, F. A. et al. Paga: graph abstraction reconciles clustering with trajectory inference through a topology preserving map of single cells. Genome biology 20, 1–9 (2019).

34. Rashid, S., Shah, S., Bar-Joseph, Z. & Pandya, R. Dhaka: variational autoencoder for unmasking tumor heterogeneity from single cell genomic data. bioRxiv 183863 (2018).

35. Haghverdi, L., Buettner, F. & Theis, F. J. Diffusion maps for high-dimensional single-cell analysis of differentiation data. Bioinformatics 31, 2989–2998 (2015).

36. Street, K. et al. Slingshot: cell lineage and pseudotime inference for single-cell transcriptomics. BMC genomics 19, 477 (2018).

37. Qiu, X. et al. Reversed graph embedding resolves complex single-cell trajectories. Nat. methods 14, 979 (2017).

38. Schiebinger, G. et al. Optimal-transport analysis of single-cell gene expression identifies developmental trajectories in reprogramming. Cell 176, 928–943 (2019).

39. Vander Velde, R. et al. Resistance to targeted therapies as a multifactorial, gradual adaptation to inhibitor specific selective pressures. Nat. communications 11, 1–13 (2020).

40. Qiu, W.-L. et al. Deciphering pancreatic islet β cell and α cell maturation pathways and characteristic features at the single-cell level. Cell metabolism 25, 1194–1205 (2017).

41. Zhu, D. et al. Single-cell transcriptome analysis reveals estrogen signaling coordinately augments one-carbon, polyamine, and purine synthesis in breast cancer. Cell reports 25, 2285–2298 (2018).

42. Mathys, H. et al. Temporal tracking of microglia activation in neurodegeneration at single-cell resolution. Cell reports 21, 366–380 (2017).

43. Guo, C. et al. Celltag indexing: genetic barcode-based sample multiplexing for single-cell genomics. Genome biology 20, 90 (2019).

44. Kong, W. et al. Celltagging: combinatorial indexing to simultaneously map lineage and identity at single-cell resolution. Nat. protocols 15, 750–772 (2020).

45. McInnes, L., Healy, J., Saul, N. & Grossberger, L. Umap: Uniform manifold approximation and projection. The J. Open Source Softw. 3, 861 (2018).

46. Coifman, R. R. et al. Geometric diffusions as a tool for harmonic analysis and structure definition of data: Diffusion maps. Proc. national academy sciences 102, 7426–7431 (2005).

47. Angerer, P. et al. destiny: diffusion maps for large-scale single-cell data in r. Bioinformatics 32, 1241–1243 (2016).

48. Mao, Q., Wang, L., Goodison, S. & Sun, Y. Dimensionality reduction via graph structure learning. In Proceedings of the 21th ACM SIGKDD International Conference on Knowledge Discovery and Data Mining, 765–774 (2015).

49. Mao, Q., Yang, L., Wang, L., Goodison, S. & Sun, Y. Simpleppt: A simple principal tree algorithm. In Proceedings of the 2015 SIAM International Conference on Data Mining, 792–800 (SIAM, 2015).

